# Distinct gene expression dynamics in germ line and somatic tissue during ovariole morphogenesis in *Drosophila melanogaster*

**DOI:** 10.1101/2021.04.28.441729

**Authors:** Shreeharsha Tarikere, Guillem Ylla, Cassandra G. Extavour

## Abstract

The survival and evolution of a species is a function of the number of offspring it can produce. In insects the number of eggs that an ovary can produce is a major determinant of reproductive capacity. Insect ovaries are made up of tubular egg-producing subunits called ovarioles, whose number largely determines the number of eggs that can be potentially laid. Ovariole number is directly determined by the number of cellular structures called terminal filaments, which are stacks of cells that assemble in the larval ovary. Elucidating the developmental and regulatory mechanisms of terminal filament formation is thus key to understanding the regulation of insect reproduction through ovariole number regulation. We systematically measured mRNA expression of all cells in the larval ovary at the beginning, middle and end of terminal filament formation. We also separated somatic and germ line cells during these stages and assessed their tissue-specific gene expression during larval ovary development. We found that the number of differentially expressed somatic genes is highest during late stages of terminal filament formation and includes many signaling pathways that govern ovary development. We also show that germ line tissue, in contrast, shows greater differential expression during early stages of terminal filament formation, and highly expressed germ line genes at these stages largely control cell division and DNA repair. We provide a tissue-specific and temporal transcriptomic dataset of gene expression in the developing larval ovary as a resource to study insect reproduction.

## INTRODUCTION

Healthy reproductive organs are among the most important factors that determine the fertility of an individual, and more importantly, continuity of the species itself. Reproductive fitness, including fecundity, is determined by the number of progenies an organism can produce. In insects, egg-producing subunits of ovaries are called ovarioles (BÜning 1994). In flies of the genus *Drosophila*, the number of ovarioles predicts the peak egg laying potential of the females of the species (David 1970), and is negatively correlated with egg size but positively correlated with reproductive output (Church *et al*. 2021). The number of ovarioles varies widely across insects and is in the range of 18-24 ovarioles per ovary in wild type North American populations of *Drosophila melanogaster* (Honek 1993; Markow and O’Grady 2007; Hodin 2009). In *Drosophila*, adult ovariole number is established in the larval stages through the development of a species-specific number of linear somatic cell stacks called terminal filaments (King *et al*. 1968). The genetic mechanisms governing ovary morphogenesis, which includes the process of regulation of terminal filament number and assembly during larval ovary development, remain poorly understood.

Ovary morphogenesis is orchestrated by interactions of the cell types of somatic and germ line tissues. Somatic ovarian tissue is principally made up of five cell types -sheath cells, swarm cells, terminal filaments, cap cells, and intermingled cells. The anterior most cells of the ovary are the sheath cells, and a sub-population of these apically positioned cells undergo two cell migration events during larval ovary development. First, a population of sheath cells called swarm cells migrates from the anterior to the posterior of the ovary to form the basal region in the mid third larval instar stage (Couderc *et al*. 2002; Green II and Extavour 2012). Secondly, in the late third instar and early pupal stages, sheath cells migrate from the apical to the basal region, traversing in between terminal filaments (King *et al*. 1968). These sheath cells lay down basement membrane in their path, which encapsulates developing ovarioles (King 1970).

Terminal filaments are stacks of cells located just below the sheath cells in the anterior larval ovary. They are formed by a process of progressive intercalation of flattened cells into stacks, and stack formation occurs in a “wave” that proceeds from the medial to the lateral side in the larval ovary (**Figure 1A**) (Sahut-Barnola *et al*. 1995). Morphogenesis in larval ovary and the mechanisms controlling the process are not completely understood.

**Figure 1:**
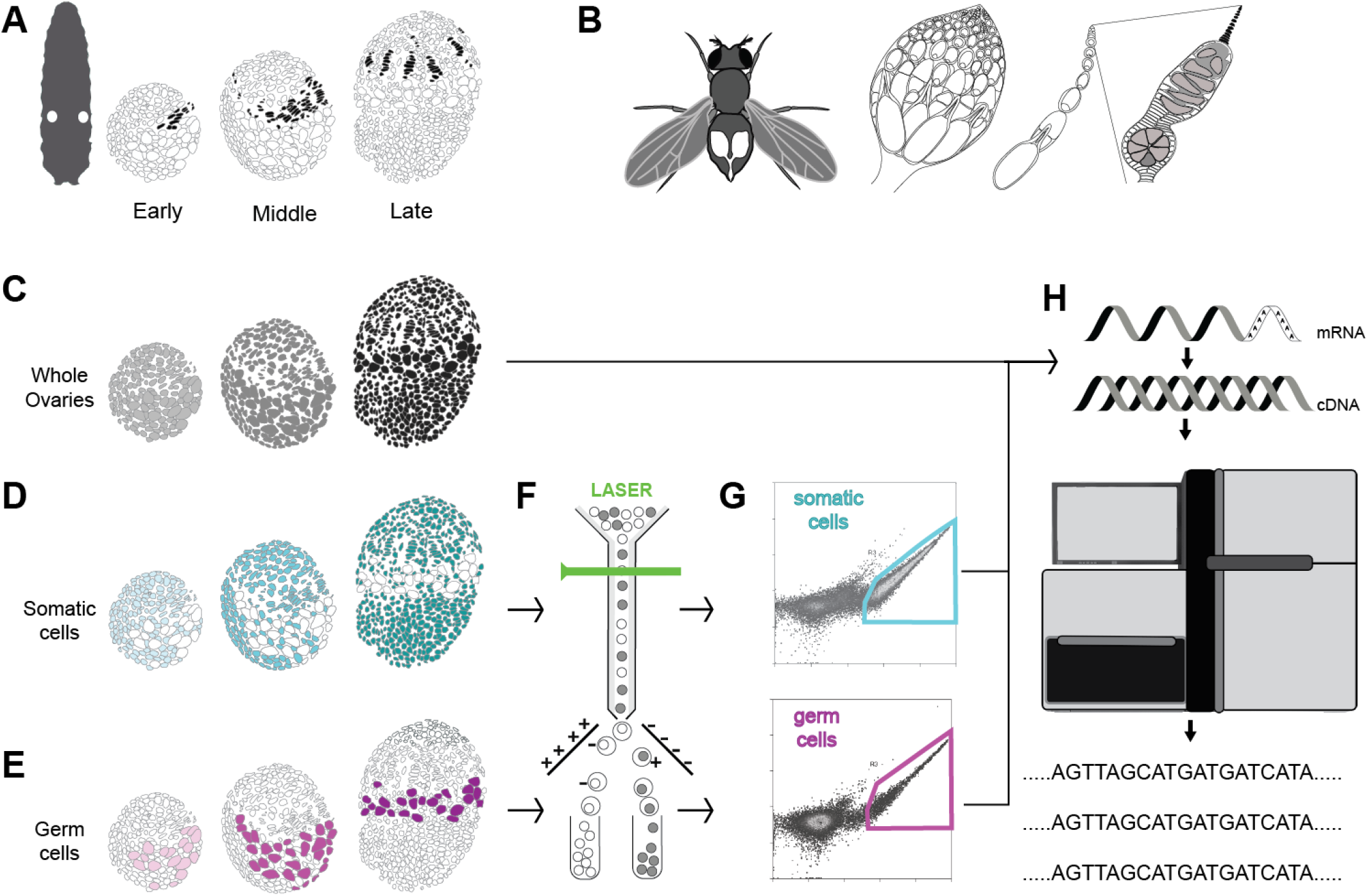
Experimental scheme for generating stage-specific transcriptomes of germ cells and somatic cells of larval ovaries during terminal filament formation. **A)** Location of the larval ovaries (white circles within the larva), and illustration of larval ovary development divided into three stages during terminal filament formation (colored in black). **B)** Left to right: location of the ovaries in an adult female abdomen; a single adult ovary containing multiple ovarioles; an individual ovariole; anterior tip of an ovariole enlarged to show the germarium and terminal filament (black) at the tip. **C)** Representation of the three stages of whole larval ovaries chosen for library preparation and sequencing (yellow: early stage, green: mid, blue: late). **D)** Somatic cells and **E)** germ cells from developing ovaries at the three chosen stages were labelled with GFP using tissue-specific GAL4 lines and **F)** GFP-positive cells were separated using FACS. **G)** Schematics of representative plot layouts of somatic and germ line tissue separation using FACS. Y axis: autofluorescence, 488-576/21 Height Log; X axis: GFP fluorescence intensity, 488-513/26 Height Log (see Supplementary Figure S2 for actual data plots). **H)** Separated cells or whole ovaries were processed for mRNA extraction and cDNA library preparation followed by high throughput sequencing.

The genes *bric á brac 1 (bab1), bric á brac 2 (bab2)* and *engrailed (en)* are expressed in the terminal filaments and essential for terminal filament cell differentiation and terminal filament assembly (Godt and Laski 1995; Sahut-Barnola *et al*. 1995; Couderc *et al*. 2002; Bolívar *et al*. 2006). We previously showed that the Hippo signaling pathway controls the regulation of cell proliferation in somatic cells, thereby affecting the number of terminal filaments and their constituent terminal filament cells (Sarikaya and Extavour 2015). During early terminal filament formation, Actin and Armadillo (arm) proteins deposited in the region between terminal filaments make a scaffold to flatten and intercalate terminal filament cells (Godt and Laski 1995; Sahut-Barnola *et al*. 1995; Chen *et al*. 2001). Expression of the protein cofilin (*twinstar*) is required in terminal filament and apical cells for actin-based change in cell shape, and loss of cofilin causes a reduction in terminal filament and apical cell numbers (Chen *et al*. 2001).

Normal growth of an ovary depends on the homeostatic of proliferation of the somatic and germ line tissues (Gilboa and Lehmann 2006; Gilboa 2015). This balance between somatic and germ line tissue populations is achieved by regulation of proliferation, differentiation and apoptosis of stem cell populations of somatic and germ cell lineages (Sahut-Barnola *et al*. 1995; Sahut-Barnola *et al*. 1996). Somatic cells called intermingled cells interact with the germ cells and control their proliferation (Li *et al*. 2003; Gilboa and Lehmann 2006; Sarikaya and Extavour 2015; Lai *et al*. 2017; Panchal *et al*. 2017; Li *et al*. 2019). *Notch, hedgehog, Mitogen Activated Protein Kinase (MAPK)* and *Epidermal growth factor receptor* (*EGFR)* signaling pathways, as well as the transcription factor *traffic jam* maintain the germ line stem cell niche (Besse *et al*. 2005; Song *et al*. 2007; Matsuoka *et al*. 2013; Sarikaya and Extavour 2015), which is established at the posterior of each terminal filament.

Recent work by Slaidina and colleagues used single-cell transcriptomics to describe the gene expression profiles of the various cell types of the late third instar larval ovary (Slaidina *et al*. 2020). They sub-divided terminal filament cells into anterior or posterior cell types, and sheath cells into migratory or non-migratory cell types, based on gene expression patterns of the single cell sub-populations. (Slaidina *et al*. 2020). While this study examined a single time point of ovary development, given that ovary morphogenesis is a temporal process, we hypothesize that changes in gene expression patterns over the course of development may be important to regulate morphogenesis. Thus, a gene expression study across the developing stages of larval ovary would advance our understanding of the transcriptomic regulation of ovarian morphogenesis.

Although all major conserved animal signaling pathways are known to be involved in ovarian morphogenesis (Twombly *et al*. 1996; Cohen *et al*. 2002; Huang *et al*. 2005; Song *et al*. 2007; Gancz and Gilboa 2013; Green and Extavour 2014; Sarikaya and Extavour 2015; Kumar *et al*. 2020), a systematic gene expression profile of a developing ovary is lacking. Such system-wide gene expression data for the ovary throughout terminal filament morphogenesis, including the potentially distinct transcriptional profiles of germ cells and somatic cells, could shed light on the processes involved in the maintenance of cell types necessary to shape the ovary and control the number of ovarioles.

To this end, we measured gene expression during the development of the larval ovary by systematically staging and sequencing mRNA from whole ovaries before, during and after terminal filament formation. Furthermore, we separated somatic and germ line tissue types at each of these stages to analyze tissue-specific gene expression. We compared the gene expression profiles across tissues and also across stages of ovary development. We then employed functional enrichment analysis to determine the different biological functions active in the three larval developmental stages and two tissue types that could yield information on ovary morphogenesis. This dataset is an important temporal and tissue specific gene expression resource for the insect developmental biology community to understand early ovary development.

## RESULTS

### Staging larval ovary development during terminal filament formation

We divided the developing *Drosophila* larval ovary into three stages during terminal filament formation and used RNA-seq to quantify gene expression at these stages (**Figure 1A**). First, we considered an early stage of terminal filament formation at the early third instar larva (72 hours After Egg Laying, 72h AEL), when terminal filament assembly is initiating (Godt and Laski 1995) (**Figure 1A-i)**. Second, we assigned the middle (mid) stage (96h AEL) as 24 hours after the early stage, at the midway point of terminal filament assembly (Godt and Laski 1995) (**Figure 1A-ii**). Third, the late stage (120h AEL) was defined as the time point of white pupa formation (when the larvae become immobile at the larval to pupal transition (Ashburner *et al*. 2005)), which occurs 24 hours after the middle stage (**Figure 1A-iii**). At the white pupa stage, terminal filament assembly is complete and the number of terminal filaments reflects the number of adult ovarioles (Hodin and Riddiford 2000) (**Figure 1B**).

We dissected these three stages of developing ovaries from larvae obtained from synchronized eggs and sequenced the transcripts present at each stage from pools of 30-100 ovaries **(Supplementary Table S1)**. We aligned reads to the *Drosophila melanogaster* genome (FlyBase v6.36), which yielded between 88.49% and 98.06% of reads aligned per sample (**Supplementary Figure S1, Supplementary Table S1**). Clustering analysis based on the variance-stabilizing transformation (VST) of the gene counts of each sample confirmed that the three biological replicates of each stage clustered together, and that the three stages were well separated, as reflected by the principal component analysis (PCA) and the dendrogram of the hierarchical analysis (**Figure 2A-B**). Furthermore, the dendrogram visualization of the hierarchical clustering results revealed that the mid stage was more similar in expression profile to the early stage than to the late stage. This indicates a more pronounced transcriptomic change at the transition from mid to late, than from early to mid, despite the fact that the same chronological amount of time had elapsed between each stage.

**Figure 2:**
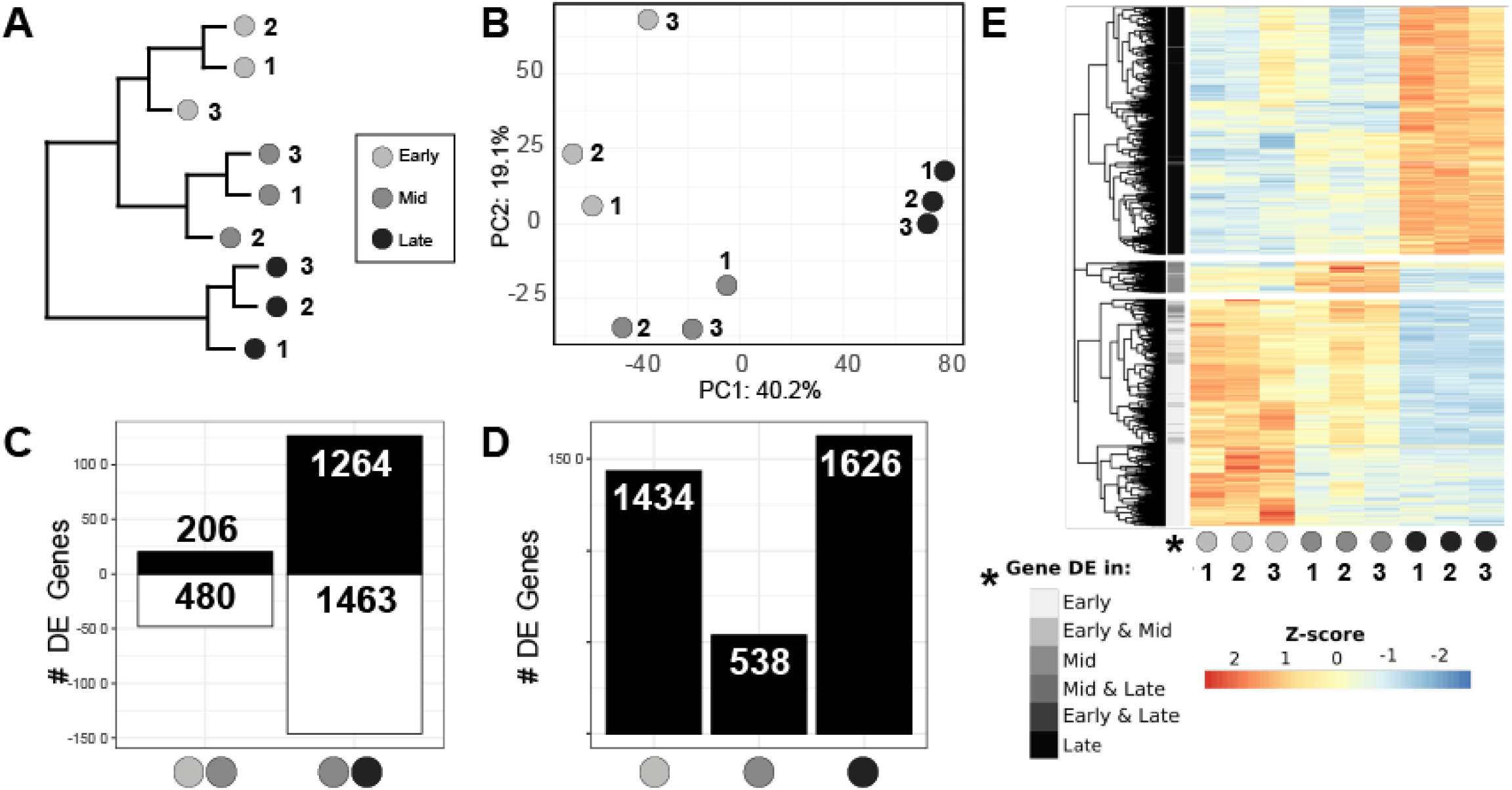
Whole ovary RNA-seq dataset overview. **A)** hierarchical clustering dendrogram and **B)** PCA of the whole ovary RNA-seq dataset, both showing that biological replicates are similar to each other and that early and mid-stages are more similar to each other than either of them is to late stage. **C)** Number of differentially expressed genes between early and mid stages, and between mid and late stages (adjusted p-value<0.01; black: upregulated genes; white: downregulated genes). **D)** Number of significantly upregulated stage-specific genes (adjusted p-value<0.01). **E)** Heatmap showing the expression of all the stage-specific upregulated genes as a row-wise z-score. Genes are clustered hierarchically and separated into three groups using the function “cutree”, and greyscale row labels immediately to the right of the tree are colored based on the stage in which the gene was detected to be significantly upregulated (x axis categories).

### Differential gene expression analysis of whole ovary samples at different stages

We analyzed the transcriptional differences between each stage and the successive one, thus performing a differential expression analysis comparing early to mid and mid to late transitions, using DESeq2 (Love *et al*. 2014) with a threshold of p<0.01 (see Methods). We found a significantly higher number of genes differentially expressed in the mid to late transition (2,727 genes), than in the early to mid transition (685) (**Figure 2C, Supplementary Table S2**). Interestingly, from early to mid stages twice as many genes were down-regulated (480) as upregulated (206), while from mid to late stages approximately the same proportion of genes were upregulated (1,264) and downregulated (1,463). We then identified the genes that were differentially expressed in one stage as compared to the other two stages, with the aim of revealing genes with stage-specific over- or under-expression. We found that early and late stages had many more over-expressed genes (1,434 and 1,626 respectively) than the mid stage (538) (**Figure 2D, Supplementary Table S3**). A heatmap representing the expression levels of the stage-specific overexpressed genes clearly separates the three groups of genes (**Figure 2E**). The first group in the heatmap contains the 1,478 genes that are highly expressed specifically at early stages, with less expression at mid stages and very low expression at the late stage. Another large group of 1,618 genes are highly expressed specifically at late stages, and show low expression at early and mid stages. Finally, we identified a third and smallest group of 202 genes that are highly expressed at mid stages, with some detectable expression at early stages, but little detectable expression at the late stage (**Figure 2E**). These results are consistent with our previous observation that there is a high gene expression similarity in early and mid-stages, and an increased transcriptomic change from mid to late stages.

### Separation of somatic and germ line tissues in the developing ovary

Given our ultimate interest in gene regulatory functions and dynamics during terminal filament formation, we wished to understand the predicted functions of the many differentially expressed genes across stages. We reasoned, however, that given the different developmental numbers, roles and behaviors of germ line and somatic cells in this developing organ, considering functional categories of differentially expressed genes in these whole ovary samples would be only minimally informative. We therefore designed an experimental strategy that allowed us to consider the transcriptional dynamics of the germ line and soma separately, described below.

To understand the gene expression differences between the somatic and germ line tissues of the ovary during terminal filament morphogenesis, we drove somatic and germ line tissue-specific GFP expression using the UAS-GAL4 system (Brand and Perrimon 1993), using the drivers *bab:GAL4* and *nos:GAL4* respectively (see Methods). We dissociated ovaries at the three stages described above, and isolated the GFP-positive cells using Fluorescence-Activated Cell Sorting (FACS). Cellular debris was eliminated with gate R1, non-singlets were eliminated by gate R2, and the R3 gate selected for GFP positive cells. A combination of the three gates yielded singlet GFP positive cells, minimizing the possibility of tissue contamination by undissociated cells. When similar number of ovaries were used to obtain sorted cells for somatic and germ line tissue-types, we found larger number of somatic cells as compared to germ cells as expected, indicating a successful separation of the desired tissue type. (**Supplementary Figure S2**).

With this method, we obtained tissue-specific transcriptomes of somatic and germ line tissues at the same three stages of terminal filament development used to generate the whole ovary dataset. We sequenced three biological replicates for all datasets, and retained replicates that had at least 10 million reads. The number of reads aligned to the genome ranged from 11 to 81 million. Greater than 94% of reads aligned in all datasets, with the single exception of one dataset with 88% of aligned reads **(Supplementary Table S1, Supplementary Figure S3**). The PCA analysis based on the counts normalized by variance-stabilization transformation (VST) shows a clear separation of somatic and germ cell libraries along the first principal component, suggesting a successful separation of cell types by FACS (**Figure 3A**). For the somatic samples, the three biological replicates cluster closely together (**Figure 3A-B**) while the different stages are separated from each other in the second principal component. The structure of the dendrogram for the somatic samples resembles that of the whole ovary, in which early and mid-stages are closer to each other than either is to the late stage. As for the germ cell libraries, unlike the biological replicates of the early and late stages, the mid-stage replicates do not cluster together. A possible explanation is the low number of reads from the sample Mid-1 (**Supplementary Table S1**).

**Figure 3:**
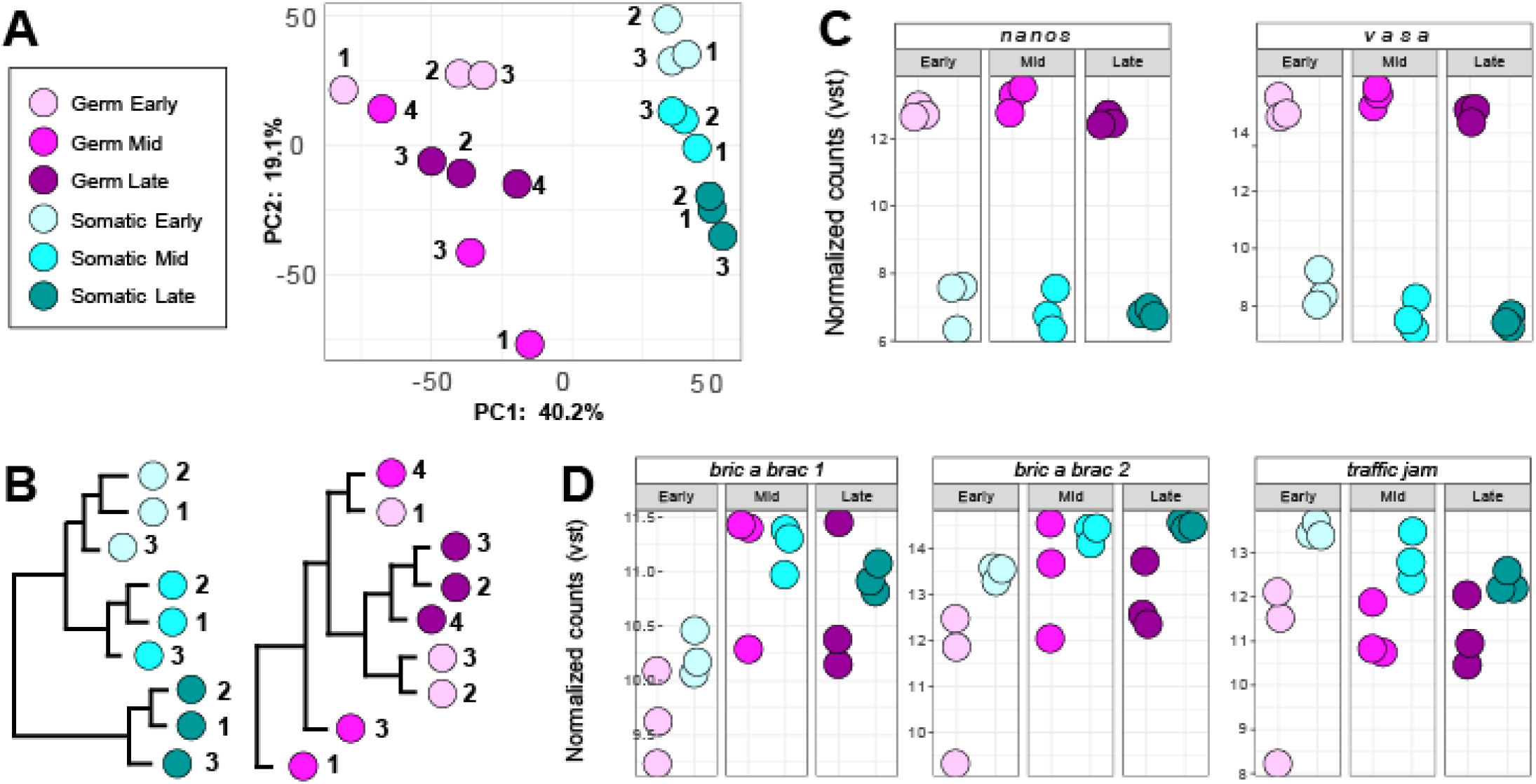
Cell type-specific RNA-seq dataset concordance and positive controls. **A)** PCA Plot and **B)** hierarchical clustering dendrogram of germ cell and somatic cell RNA-seq libraries. Expression in normalized counts by variance stabilization transformation (VST) in each of the cell-type-specific RNA-seq libraries of **C)** known germ cell markers *nanos* and *vasa*, and **D)** known terminal filament markers *bric a brac 1, bric a brac 2*, and *traffic jam*.

To further assess the successful separation of somatic and germ cells, we checked the expression of well-known tissue-type-specific markers. The genes *nanos* and *vasa* are two genes known to be specifically expressed in germ cells in the ovary (Schupbach and Wieschaus 1986; Lehmann and Nusslein-Volhard 1991). Both genes show higher expression in the germ cell libraries than in the somatic cell libraries (mean log_2_(Fold Change) of 8.37 for *nanos*, and 8.23 for *vasa*) (**Figure 3C**), confirming that the preparation and sequencing of the germ cell libraries successfully captured the germ cells and their RNAs, and suggesting that germ cells were not present (or present only at very low levels) in the somatic cell libraries. *bab1, bab2*, and *tj* are considered somatic gene markers (Sahut-Barnola *et al*. 1995; Couderc *et al*. 2002) These three somatic markers display higher expression levels in our somatic libraries than in the germ cell libraries at each stage (mean log_2_(Fold Change) -0.31 for *bab1*, -1.3 for *bab2*, and -1.7 for *tj*) (**Figure 3D**). However, in four of the 18 libraries, either *bab1* or *bab2* (but not *tj*) showed higher expression levels in a specific germ cell library than in the somatic libraries. These specific cases were as follows: (1) one early stage germ cell replicate had higher *bab1* levels than one of the early somatic replicates; (2) two mid stage germ cell replicates had higher *bab1* levels than the somatic replicates; (3) one late stage germ cell replicate had higher *bab1* levels than the somatic replicates; (4) one mid stage germ cell replicate had higher *bab2* levels than the somatic replicates. This could indicate that some somatic cells might have been included in these particular germ cell libraries. Nonetheless, despite this putative small amount of contamination, we can clearly differentiate both tissue types based on their expression profiles as shown in the PCA (**Figure 3A**), suggesting that we captured the transcriptional differences between cell types (**Figure 3A**) sufficiently to allow us to achieve our goal of successfully retrieving the genes that are highly and differentially expressed in each of these two tissues.

### Differential expression analysis of somatic and germ line tissues across all stages

The differential expression analysis between the somatic and germ line tissues across all three stages revealed 1,880 genes significantly upregulated (adjusted p-value<0.01) in germ cells and 1,585 genes significantly upregulated in the somatic cells (**Figure 4A; Supplementary Table S4**).

**Figure 4:**
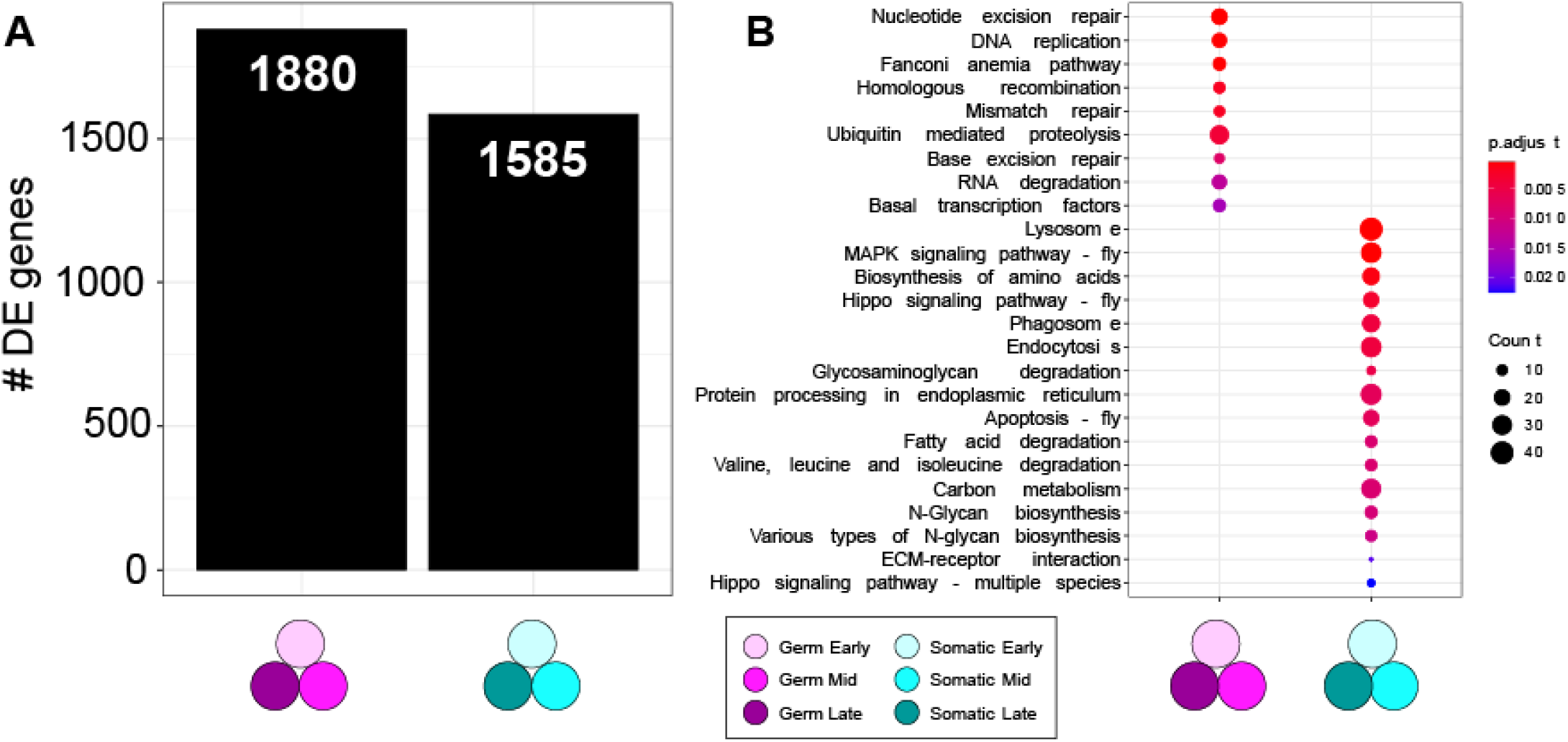
Transcriptomic differences between germ cells and somatic cells. **A)** Number of significantly upregulated genes (adjusted p-value<0.01) in germ cells and somatic cells. **B)** Significantly enriched KEGG pathways (adjusted p-value<0.05) within the upregulated genes of each cell type. The circle size is proportional to the number of differentially expressed genes that the indicated KEGG pathway contains, and the color gradient indicates the p-value.

Among the 20 most significant genes (with the lowest adjusted p-value) overexpressed in germ cells relative to somatic cells, we detected known germ line-specific genes including piRNA biogenesis genes *Argonaute3 (AGO3), krimper (krimp)*, and *tejas (tej)*, along with *Aubergine (aub)*, (Brennecke *et al*. 2007; Olivieri *et al*. 2010; Patil and Kai 2010; Sato *et al*. 2015), *sisters unbound (sunn)* (Krishnan *et al*. 2014), *benign gonial cell neoplasm (bgcn)* (Ohlstein *et al*. 2000), and uncharacterized genes including *CG32814* and *CG12851* on the chromosome 2R. As for the somatic cells, the most significantly overexpressed gene relative to the germ cells is the cytochrome gene *Cyp4p2*, whose role is unknown in the ovary, followed by cytochrome *Cyp4p1* and the uncharacterized genes CG32581 and CG42329. Some genes known to play roles in the ovary were also among this group, including the regulator of the niche cells and ecdysone receptor *Taiman (tai)* (KÖnig *et al*. 2011), and the regulator of vitellogenesis *apterous (ap)* (Gavin and Williamson 1976).

### Temporally dynamic expression of genes previously studied in somatic ovary development

We explored the expression dynamics of some of the previously studied genes expressed in the *Drosophila* ovary. To our knowledge, temporal gene expression studies in the larval ovary for many these genes have not yet been conducted.

First, we considered the temporal expression patterns of some adhesion proteins known to play a role in ovary development. *RanBPM* is an adhesion linker protein expressed in the germ line niche in the adult ovary (Dansereau and Lasko 2008). In our dataset we see opposing trends of expression levels in somatic and germ line tissues, such that in germ line tissue *RanBPM* expression decreases progressively from early to mid to late stages, while in the somatic tissue it increases from early to mid to late stages (**Supplementary Figure S4A**). Cofilin (encoded by the gene *twinstar*) is an adhesion protein required for terminal filament cell rearrangement during terminal filament morphogenesis, as well as for adult border cell migration (Chen *et al*. 2001). Cofilin shows similar germ line and somatic cell expression trends, with higher levels at early stages that decrease progressively at mid and late stages (**Supplementary Figure S4B**).

We then looked at temporal expression of *RhoGEF64C* and *Wnt4*, genes involved in cell motility. RhoGEF64C is a small apically localized RhoGTPase that regulates cell shape and migration in the ovary (Simoes *et al*. 2006). In our datasets we found *RhoGEF64C* expressed at higher levels in early and late stage somatic cells than at mid stages (**Supplementary Figure S4C**). *Wnt4* is involved in cell motility during ovarian morphogenesis (Cohen *et al*. 2002) and is expressed in the posterior terminal filaments and other somatic cell types of the third instar larval ovary (Slaidina *et al*. 2020). We found *Wnt4* to be expressed in lower levels in early and middle stages while the expression increases significantly in the late stage (**Supplementary Figure S4D**).

We also examined the temporal expression dynamics of a number of terminal filament cell-type-specific genes previously identified in a single cell sequencing study of the late third larval instar ovary (Slaidina *et al*. 2020). For example, *Diuretic hormone 44 receptor 2 (Dh442)* was identified as highly expressed in terminal filament cells (Slaidina *et al*. 2020). In our datasets, we observed a significant increase in expression levels only at the late stage relative to early and mid-stage expression levels (adjusted p-value 2.04×10^−11^) (**Supplementary Figure S4E**). Additional genes known to function in terminal filaments are *engrailed* and *patched* (Forbes *et al*. 1996; Besse *et al*. 2005; Bolívar *et al*. 2006). In our datasets we observed *engrailed* expressed at lowest levels at the early stage, showing a progressive increase in expression levels from mid to late stages. *patched* showed a similar progressive increase across stages, with a significant increase from early to mid-stage (**Supplementary Figure S4F-G**). Finally, we considered members of the Fibroblast Growth Factor (FGF) signaling pathway, which controls sheath cell proliferation in the pupal ovary (Irizarry and Stathopoulos 2015). Three key genes of this pathway, the FGF ligand *thisbe*, the FGF scaffolding protein *stumps* and the upstream FGF signaling activator *heartless*, show significantly higher differential expression levels at early to mid-stage than at mid to late stages (**Supplementary Figure S4H-J**). These temporal profiles add to our understanding of the roles of these genes in ovarian morphogenesis by suggesting distinct putative critical regulatory periods for different genetic pathways.

### Functional enrichment analysis of differentially expressed genes in somatic and germ line tissues across all stages

To gain insight into the general functional categories of genes likely involved in ovarian germ cell and somatic behaviors during terminal filament development, we performed a gene ontology (GO) enrichment analysis of the biological processes of differentially expressed genes across cell types and developmental stages (Ashburner *et al*. 2000). We found 31 level four GO-terms enriched (adjusted p-value<0.05) within the upregulated genes in germ cells, and 188 level four GO-terms enriched in the upregulated genes in somatic cells (**Supplementary Figure S5**). This analysis highlighted clear differences in the biological functions performed by the genes expressed in each tissue. The GO-terms enriched in the germ cells are primarily related to meiotic processes (9/31 contain the words “meiosis” or “meiotic”), chromosome stability (6/31 contain the words “chromosome” or “karyosome”) and cell cycle (12/31 contain “cell cycle”). In contrast, the GO-terms enriched in the somatic cells are principally related to cellular response (21/188 contain “response”), development (18/188), growth (16/188), morphogenesis (10/188) cell migration (6/188 contain the word “migration”) and signaling pathways (6/188).

To complement this GO enrichment analysis, we performed a KEGG pathway enrichment analysis on the same cell-type-specific overexpressed genes. The KEGG pathway database is a manually curated database of molecular interactions used to study enrichment of genetic regulatory pathways in gene lists (Kanehisa and Goto 2000). With this analysis, we identified nine KEGG pathways significantly enriched in the germ cells, and 16 significantly enriched pathways in the somatic cells (adjusted p-value<0.05) (**Figure 4B**). The KEGG pathways enriched in the germ cells are generally related to meiosis and genome protection, while upregulated genes in the somatic cells are enriched for pathways involved in cell proliferation and cell death, including the previously identified *Hippo* (Barry and Camargo 2013; Sarikaya and Extavour 2015) and *MAPK* (Shaul and Seger 2007) signaling pathways.

### Stage- and tissue-specific differential gene expression analysis

To obtain a finer-grained view of the dynamic regulation of terminal filament development, we also performed differential expression analysis between the somatic and germ line tissue types at each of the three stages. In the somatic cells the number of differentially expressed genes between the early and mid-stages (867 genes) is lower than between the mid and late stages (1,404 genes) (**Figure 5A; Supplementary Table S6**). To identify genes with stage-specific upregulation, we compared each stage to the other two stages. We identified a higher number of stage-specific upregulated genes in early (1,227) and late stages (139) than at mid (1,409) (**Figure 5B; Supplementary Table S7**).

**Figure 5:**
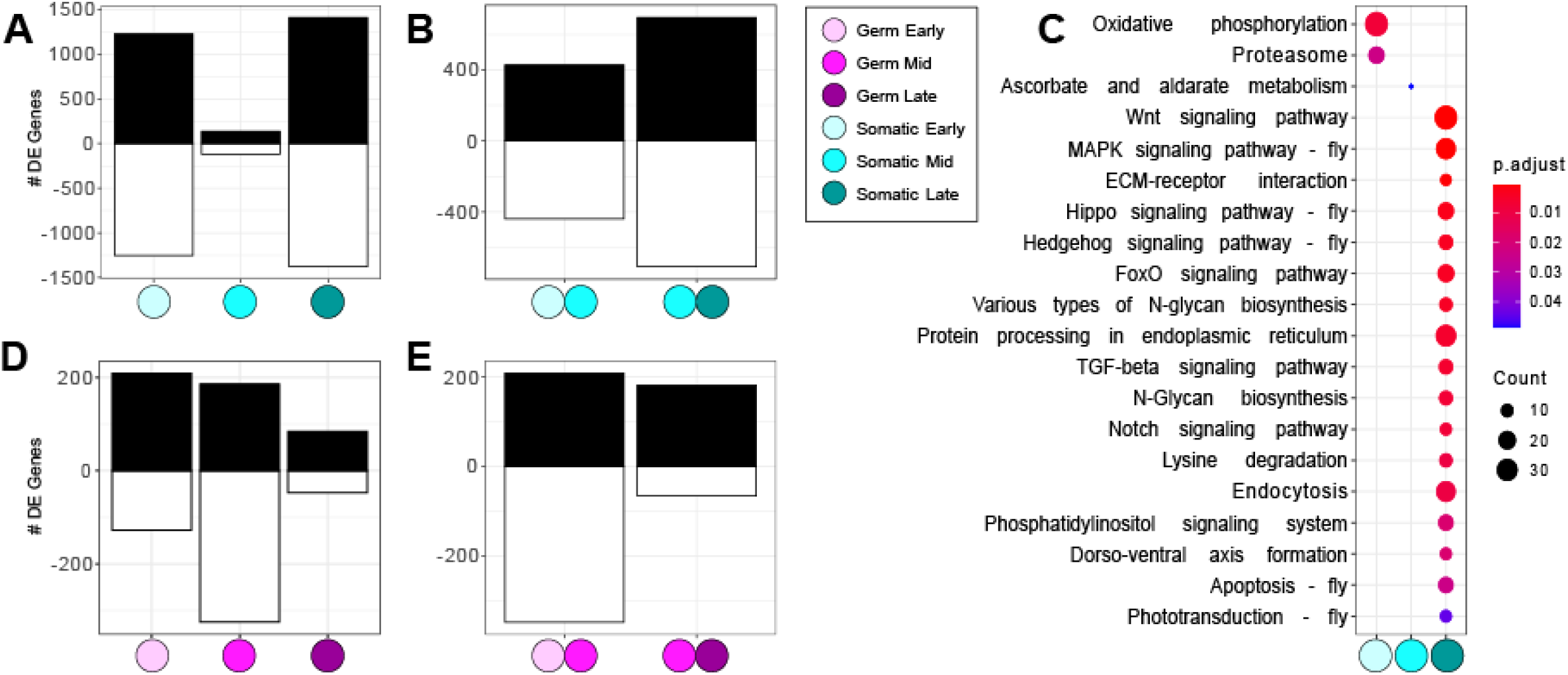
Cell type-specific differential expression analysis. **A)** Number of differentially expressed genes (adjusted p-value<0.01) upregulated (black) and downregulated (white) in somatic cells at each stage compared to the two other stages. Number of differentially expressed genes (adjusted p-value<0.01) upregulated (black) and downregulated (white) in somatic cells at each examined stage. **C)** Significantly enriched (adjusted p-value<0.05) KEGG pathways within the upregulated genes at each somatic stage. Circle size is proportional to the number of differentially expressed genes it contains, and the color gradient indicates the p-value. **D)** Number of differentially expressed genes (adjusted p-value<0.01) upregulated (black) and downregulated (white) in germ cells at each stage compared to the two other stages. **E)** Number of differentially expressed genes (adjusted p-value<0.01) upregulated (black) and downregulated (white) in germ cells at each examined stage.

The germ cells, in general, display fewer differentially expressed genes between stages than the somatic cells. From early to mid-stages there are twice as many differentially expressed genes (557 genes) as from mid to late stages (248 genes) (**Figure 5D; Supplementary Table S8**). In terms of stage-specific upregulated genes, the highest number of such genes are found at early stages (209), followed by mid (186), and late (84) stages (**Figure 5E; Supplementary Table S9**).

To explore the functions of the stage-specific upregulated genes in each tissue type, we performed a GO analysis of biological functions and KEGG pathway enrichment analysis on these six sets of genes (upregulated at early, mid, and late stages in germ and somatic cells). The GO enrichment analysis of the genes differentially expressed in somatic cells over time (**Supplementary Table S6**) revealed that four key biological processes are consistent throughout all three stages including the mid stage, which has the smallest number of differentially expressed genes across stages. Specifically, these are the GO terms taxis, cell growth, actin filament-based process and cell adhesion. At early and late stages, we additionally observe many key biological processes related to morphogenesis in the somatic cells, including cell proliferation, differentiation and migration. Considering gene expression levels specific to each stage, we identified 1,227, 139, and 1,409 upregulated genes at early, mid, and late stages respectively. The upregulated genes at early and late stages were enriched for 97 and 764 GO-terms of biological process, while none were enriched in the mid-stage (**Supplementary Figure S6**). As for the KEGG pathway enrichment analysis, there were two enriched pathways at early stages, one at mid-stage and 17 in late stages (**Figure 5C**). This analysis allowed us to pinpoint the stage(s) at which specific pathways were enriched in somatic cells relative to germ cells, which included Apoptosis, *Hippo* signaling, and *MAPK* signaling. In addition, we detected some signaling pathways enriched only in somatic cells at late stages, such as the *Hedgehog, FoxO*, and *Notch* pathways (**Figure 5C**).

Given the known role of the Hippo pathway in cell proliferation (Barry and Camargo 2013), and specifically in terminal filament cell and terminal filament number regulation (Sarikaya and Extavour 2015), we proceeded to analyze the expression patterns of the genes belonging to the core Hippo signaling pathway. We found that most Hippo pathway core genes display increasing expression levels from early to mid to late stages, with the exception of the expression of the core gene *Rae1* which progressively decreases in expression level from early to late stages (**Supplementary Figure S8**).

In the germ cells, across stages we find fewer processes directly involved in development and morphogenesis with gene ontology categories belonging to meiosis and cell cycle (**Supplementary Figure S6**). Among the 209, 186, and 84 upregulated genes in germ cells at early, mid, and late stages respectively, only one KEGG pathway (Ribosome,) and one biological process GO-term (cytoplasmic translation) of were found significantly enriched at early-stages. No such enrichment was detected at mid-stages, and three KEGG pathways were enriched at late stages (**Supplementary Figure S7**).

### Uncharacterized genes

The detection of uncharacterized genes among the top differentially expressed genes in germ cells drove us to ask if there were any differences in the proportion of uncharacterized genes in each set of differentially expressed genes. We found that in the genes significantly upregulated in somatic cells compared to germ cells, 29.15% are categorized as “uncharacterized proteins” in FlyBase (Larkin *et al*. 2021), while within the significantly upregulated genes in germ cells, the proportion of uncharacterized genes was 39.10%. Within the stage-specific upregulated genes, the proportion of uncharacterized genes remained constant (between 28.96% and 29.63%) in somatic cells, while in germ cells it increased from 29.08% in early stages, to 34.83% in mid stages, and to 37.40% in late stages (**Supplementary Figure S9)**.

### Expression of cell type-specific markers

A previous single cell RNA-sequencing dataset of the late third stage larval ovary () Slaidina *et al*. (2020) identified transcriptional profile clusters interpreted as indicative of cell types, and suggested gene markers associated with each cell type. To determine whether the cell types identified at this late stage might also be present at earlier developmental stages than that previously assessed, we examined the expression levels of those suggested marker across our datasets. As expected, the majority of the germ cell markers are highly expressed in our germ cell libraries and expressed only at low levels in the somatic cells (**Supplementary Figure S10**). Among the somatic markers detected in our somatic tissue libraries, we do not observe any particular temporal expression pattern specific to a given somatic cell type. Nevertheless, we clearly distinguish two groups of somatic markers (**Supplementary Figure S11**). One group is composed of somatic markers whose expression levels are highest at early and mid-stages, and decay at the late stage, and a larger group of makers that are less strongly expressed at early stages, show increased expression at the mid stage, and show highest expression at late stages. By contrast, the germ cell markers detected in our germ cell libraries do not display any clear temporal expression pattern. Instead, most of these genes were expressed at similar levels across the three studied stages (**Supplementary Figure S12**). This is consistent with our previous observations that the germ line dataset is not enriched for any signaling pathway directly implicated in development during these three times points as the somatic cells do (Supplementary Figure S6).

## DISCUSSION

### Temporal gene expression during ovary morphogenesis

We systematically staged and sequenced entire larval ovaries to generate a gene expression dataset during terminal filament formation. We then separated somatic and germ line tissues during these stages and generated tissue-specific transcriptomes. While the development of the *Drosophila* ovary has been studied for the last several decades, and progress has been made on identifying the roles of some signaling pathways in its morphogenesis (Cohen *et al*. 2002; Besse *et al*. 2005; Gilboa and Lehmann 2006; Gancz *et al*. 2011; Gancz and Gilboa 2013; Matsuoka *et al*. 2013; Gilboa 2015; Irizarry and Stathopoulos 2015; Lengil *et al*. 2015; Mendes and Mirth 2016; Panchal *et al*. 2017) to our knowledge, there are no publicly available transcriptomes of developing larval ovaries of *Drosophila*. Recent articles have reported single cell RNA-sequencing for *Drosophila* ovaries, focusing either on a single larval time point or on adult ovaries (Jevitt *et al*. 2020; Rust *et al*. 2020; Slaidina *et al*. 2020; Slaidina *et al*. 2021). Our stage and tissue-type specific data thus represent a valuable complementary transcriptomic resource on the morphogenesis of the larval ovaries of *Drosophila*, a complex process that ultimately influences reproductive capacity.

In the whole ovary dataset, the increased differential gene expression at the mid-late transition and at the late stages, enriched the expression of genes in key signaling pathways that are necessary in ovary development (**Figure 2C-D**). Signaling pathways found exclusively enriched at the late stages suggest that morphogenetic processes of the larval ovary at these stages operate through these key pathways.

### Separating somatic and germ line tissue

FACS-based tissue-type separation coupled with RNA-seq proved to be a successful way to obtain transcriptomes of somatic and germ line cells during larval ovary development. The overall expression profile of the somatic cells across the three studied stages is similar to the profile of the whole ovary dataset in terms of the proportion of differentially expressed genes across stages(**Figures 2D, 5B**) This may be because since the number of somatic cells in the whole ovary is higher than that of the germ cells at all stages (**Supplementary Figure S2**), the whole ovary gene expression profile is likely dominated by somatic cell expression. Only upon physically separating the germ cells from the somatic cells, could we observe a different gene expression pattern in the germ line. Germ cells showed the highest number of differentially expressed genes at early stages, and the lowest numbers at late stages (**Figure 5E**). In contrast, the highest number of differentially expressed genes of somatic stages occurs at the late stages, and the lowest number at the mid stage (**Figure 5B**).

The results of the functional enrichment analyses in somatic and germ line tissue reveal distinct functions and pathways likely operate in these tissues during larval ovary development. Germ cells may be especially sensitive to DNA damage given their role in propagating genetic material, which we speculate may explain the enrichment of processes related to nucleotide replication, recombination and repair in our analysis (Figures 4, S5). Similarly, we observed many genes of the piRNA pathway (e.g., *AGO3, aub, krimp, tej*), which protect the genome from transposable elements (**Supplementary Table S6**) (Sato and Siomi 2020) among the top significantly enriched genes in germ cells. On the other hand, the somatic tissue is enriched for different signaling pathways including Hippo, MAPK, and apoptosis (**Figures 5C, S6**), which are known to play a role in either larval or adult ovary morphogenesis (Lynch *et al*. 2010; Khammari *et al*. 2011; Elshaer and Piulachs 2015; Sarikaya and Extavour 2015).

Detection of a higher number of uncharacterized genes in germ line tissue datasets than in the somatic tissue (**Supplementary Figure S9**), suggest that our understanding of the genetic regulation of the germ line in developing ovaries is still incomplete. Datasets like the one herein provided help to identify new genes that could be important for ovary morphogenesis.

### Cell adhesion and migration during ovary morphogenesis

We assessed the temporal dynamics of genes expressed in specific cell types during development to serve as generators of new hypotheses to understand the role of genes and pathways during morphogenesis. *RhoGEF64C* is a RhoGTPase with some role in regulating control cell shape changes that lead to epithelial cell invagination (Simoes *et al*. 2006; Toret and Le Bivic 2021). In a genome-wide association study on ovariole number phenotypes in natural populations of *Drosophila, RhoGEF64C* driven in somatic tissue had a significant effect on adult ovariole number (Lobell *et al*. 2017). The significant increase in expression of *RhoGEF64C* we observed in early and late stages (Figure S4C) suggests its role in somatic cell shape and migration in both early and late stages. We see the GO processes of adhesion and migration enriched in somatic cells but not in germ cells (**Supplementary Figure S5**), which could mean that these morphogenetic processes in germ cells are minimal compared to somatic cells.

GO-terms related to cell adhesion, motility and taxis were enriched in all three stages in somatic cells (**Supplementary Figure S6**). Previous studies have shown signaling pathways involved in ovary development to affect cell adhesion and migration processes (Cohen *et al*. 2002; Li *et al*. 2003; Besse *et al*. 2005; Lai *et al*. 2017). Migratory events in mid to late stages of the larval ovary have been described for two ovarian cell types, swarm cells and sheath cells (Sahut-Barnola *et al*. 1995; Sahut-Barnola *et al*. 1996; Green II and Extavour 2012; Slaidina *et al*. 2020). Migrating sheath cells in the late third instar larvae lay the basement membrane in between terminal filaments to form ovarioles (King *et al*. 1968).

The FGF signaling pathway supports terminal filament cell differentiation in the early larval stages through *thisbe (ths)* and upstream activator *heartless (htl)*, and also controls sheath cell proliferation in late larval and pupal stages (Irizarry and Stathopoulos 2015). In our dataset, we observe that in somatic cells these FGF pathway genes show a significant progressive upregulation from early to mid and from mid to late stages (**Supplementary Figure S4I-J**). Consistently, *ths* and *stumps* were identified as markers of a distinct migratory ovarian cell population, the sheath cells (Slaidina *et al*. 2020). *stumps* is expressed in stages corresponding to our “late” stage in the differentiating terminal filament cells and at later stages (144h AEL), also in migratory sheath cells (Irizarry and Stathopoulos 2015).

### Functional enrichment analysis and signaling pathways

Our results show that in the late stage of somatic cells there is an increase in expression of genes involved in multiple signaling pathways, including the Wnt, MAPK, Hippo, Hedgehog, FoxO, TGF and Notch pathways (**Figure 5C**). The molecular mechanisms of all these signaling pathways during larval ovary development have not yet been extensively studied, but all of them have been functionally implicated in ovariole number determination by a large-scale genetic screen (Kumar *et al*. 2020).

We previously showed that Hippo signaling pathway controls proliferation of somatic cells, which affects terminal filament number (Sarikaya and Extavour 2015). Our differential gene expression data show that members of the Hippo pathway are significantly differentially expressed in the somatic tissue (**Figure 4B; Supplementary Figure S8**), and all of its core genes except one show a progressive increase in expression levels across the three studied stages (**Supplementary Figure S8**). Loss of function mutations in Yki, an effector of the Hippo signaling pathway, cause increased growth and reduced apoptosis through an increase in the levels of the cell cycle protein Cyc E and the apoptosis inhibitor *Diap1* (Harvey *et al*. 2003; Huang *et al*. 2005). In our somatic cell datasets, we observe *Diap1* transcript levels significantly increase from early to late stages, and those of CycE increase from mid to late stages (**Supplementary Figure S13A-B**). However, the Apoptosis KEGG pathway appears significantly enriched in the somatic late stage (**Figure 5C; Supplementary Figure S6**) . Furthermore, apoptosis-related genes *Dronc* and *Dark*, which form the apoptosome (**Supplementary Figure 13C, D**) (Yuan *et al*. 2011), are also significantly upregulated in the late stage, as are the caspases *Dcp-1, Drice*, and *Dredd* (**Supplementary Figure 13C, E-G**) (Harvey *et al*. 2001). Thus, we observe both an upregulation of apoptosis and an upregulation of the apoptosis inhibition genes in late stage somatic cells. This could mean that genes controlling apoptosis both positively and negatively are acting to exert tight control of this process. Alternatively, our observations may reflect that each process is upregulated within different somatic cell types.

Cap cells and intermingled cells are somatic cells that interact with the germ cells for the maintenance of germ line stem cell niches (Li *et al*. 2003; Song *et al*. 2007). The Notch signaling pathway, enriched in the late-stage somatic dataset (**Figure 5C; Supplementary Figure S6**), is required for cap cell fate (Panchal *et al*. 2017). We observed an expression level increase in Notch pathway components at late stages, suggesting that the role of the Notch pathway in cap cell fate determination may be particularly important at mid to late stages of larval ovary development.

Components of the TGFβ pathway, enriched in late stage somatic cells in our dataset (**Figure 5C**), are known to contribute to ovarian development. These include the TGFβ component *decapentaplegic* (*dpp)*, previously documented as expressed in all larval ovarian somatic cells (Sato *et al*. 2010) and also in the larval-pupal stage cap cells and intermingled cells of the germ line stem cell niche, where it promotes proliferation and represses differentiation of primordial germ cells (Gilboa and Lehmann 2004; Matsuoka *et al*. 2013). The activin pathway, a branch of the TGFβ pathway (Pangas and Woodruff 2000), controls terminal filament cell proliferation and differentiation (Lengil *et al*. 2015). We find that the activin receptor *baboon* shows a significant expression level increase in the late stage somatic cells (adjusted p-value of 0.002602) and could indicate its role in terminal filament cell differentiation in late stage.

## Conclusions

Here we provide a dataset that explores gene expression during larval ovary development and morphogenesis, which is crucial to understand how the ovary is shaped in early stages to develop into a functional adult organ. This study advances our understanding of the process of building an ovary and regulating the morphogenesis processes. More importantly, this work offers a dataset for the developmental biology community to probe the genetic regulation of larval ovarian morphogenesis.

## MATERIALS AND METHODS

### Fly Stocks

Flies were reared at 25°C at 60% humidity with food containing yeast and in uncrowded conditions. The following two fly lines were obtained from the Bloomington Stock Center: *w[*]; P{w[+mW*.*hs]=GawB}bab1[Pgal4-2]/TM6B, Tb[1]* (abbreviated herein as *bab:GAL4*; stock number 6803), *P{w[+mC]=UAS-Dcr-2*.*D}1, w[1118]; P{w[+mC]=GAL4-nos*.*NGT}40* (abbreviated herein as *nos:GAL4*; stock number 25751). *w[1118], P[UAS Stinger]* (abbreviated herein as *UAS:Green Stinger I*, (Barolo *et al*. 2000) used for GFP expression was a gift from Dr. James Posakony (University of California, San Diego). Crosses were set with 100-200 virgin UAS females and 50-100 GAL4 males in a 180 ml bottle containing 50ml standard fly media one day prior to egg laying.

### Staging larvae

To obtain uniformly staged larvae for the experiments, a protocol was devised to collect eggs that were near-synchronously laid, from which the larvae were then collected. To obtain a desired genotype, crosses were set as described above. The cross was set at 25°C at 60% humidity and left overnight to mate. Hourly egg collections were set up on 60 mm apple juice-agar plates (9 g agar, 10 g sugar, 100 ml apple juice and 300 ml water) with a pea-sized spread of fresh yeast paste (baker’s yeast granules made into a paste in a drop of tap water). Eggs were collected hourly for eight hours. The first two collection plates were discarded to remove asynchronously laid eggs that may have been retained inside the females following fertilization. Staged first instar larvae were collected into vials 24 hours after egg collection. Larvae at 72h AEL (hours After Egg Laying) were designated as early stage, at 96h AEL as middle stage and at 120h AEL as late stage of Terminal Filament development. For a step-by-step detailed protocol see **Supplementary File 1**.

### Dissection and dissociation of larval ovary

Staged larvae were collected for dissection every hour. The head of the larva was removed with forceps and the cuticle and gut were carefully pulled with one forceps while holding the fat body with another forceps. This process left just the fat bodies in the dissection dish as long as the larvae were well fed and fattened with yeast. Ovaries located in the center of the length of each fat body were then dissected free of the fat body using an insulin syringe needle (BD 328418). Ovaries dissected clear of fat body were collected in DPBS (Thermo Fisher 14190144) and batches of 20-30 ovaries in DPBS were kept on ice until dissociation. Ovaries were harvested hourly at the appropriate times, placed on ice immediately following dissection, and maintained on ice for a maximum of four hours before dissociation and subsequent FACS processing.

Dissociation of the larval ovary required two enzymatic steps. After seven hours of dissection, batches of dissected ovaries were placed in 0.25% Trypsin solution (Thermo Fisher 25200056) for ten minutes at room temperature in the cavity of a glass spot plate (Fisher Scientific 13-748B). They were then transferred to another cavity containing 2.5 % Liberase (5 g Liberase reconstituted in 2ml nuclease free water; Sigma 5401119001) and teased apart with tungsten needles until most of the clumps were separated and left (without agitation) at room temperature for ten minutes. Using a 200µl pipette with a filter tip (pre rinsed in 1X PBS), the dissociated cells in Liberase were pipetted up and down gently ten times to uniformly mix and separate the cells. The cell suspension was then transferred to an RNA Lobind tube (Eppendorf 8077-230) and placed on a vortexer for 1 minute. Meanwhile the well was rinsed in 1.4 ml of PBS by pipetting repeatedly. This PBS was then mixed with the Liberase mixture and vortexed for another minute, and the entire sample was then was placed on ice. This sample was then taken directly to the FACS facility on ice along with an RNA Lobind collection tube containing 100-200µl Trizol (Thermo Fisher 15596026). For a step-by-step detailed protocol see **Supplementary File 1**.

### Flow Sorting GFP-positive cells

The dissociated tissue sample was sorted in a MoFlo Astrios EQ Cell sorter (Beckman Coulter) run with Summit v6.3.1 software. The dissociated cell solution was diluted and a flow rate of 200 events per second was maintained with high sorting efficiency (< 98%) during the sorting process. A scatter gate (R1) was employed to eliminate debris (**Supplementary Figure S6**) and a doublet gate (R2) was used to exclude non-singlet cells. A 488 nm emission Laser was used to excite the GFP and the collection was at 576 nm. The GFP-positive cells were designated in gate R3 and sorted directly into Trizol. The resulting cells collected in Trizol were frozen immediately by plunging the tube in liquid nitrogen and then stored at -80°C until RNA extraction. A single replicate consisted of at least 1000 cell counts pooled from FACS runs.

### RNA extraction

Flow-sorted cells were stored at -80°C were thawed at room temperature. Trizol contents were lysed with a motorized pellet pestle (Kimble 749540-0000). Zymo RNA Micro-Prep kit (Zymo Research R2060) was used to isolate RNA from the Trizol preparations. Equal amounts of molecular grade ethanol (Sigma E7023) were added to Trizol and mixed well with a pellet pestle, then pipetted onto a spin column. All centrifugation steps were done at 10,000g for one minute at room temperature. The column was washed with 400µl Zymo RNA wash buffer and then treated with Zymo DNase (6U/µl) for 15 minutes at room temperature. The column was then washed twice with 400µl Zymo RNA Pre-wash buffer and once with Zymo RNA wash-buffer. The RNA was eluted from the column in 55 µl of Nuclease-free water (Thermo Fisher 10977015). The RNA obtained was quantified first using a NanoDrop (Model ND1000) spectrophotometer and then using a high sensitivity kit (Thermo Fisher Q32852) on a Qubit 3.0 Fluorometer (Thermo Fisher Q33216). It was also checked for integrity on a high sensitivity tape (Agilent 5067-5579) with an electronic ladder on an Agilent Tapestation 2200 or 4200. RNA extraction from staged whole ovaries was carried out by crushing entire ovaries in Trizol and following the same protocol described above. For a step-by-step detailed protocol see **Supplementary File 1**.

### Library Preparation

cDNA libraries were prepared using the Takara Apollo library preparation kit (catalogue # 640096). Extracted RNA samples were checked for quality using Tapestation tapes. 50µl of RNA samples were pipetted into Axygen PCR 8-strip tubes (Fisher Scientific 14-222-252) and processed through PrepX protocols on the Apollo liquid handling system. mRNA was isolated using PrepX PolyA-8 protocol (Takara 640098). The mRNA samples were then processed for cDNA preparation using PrepX mRNA-8 (Takara 640096) protocol. cDNA products were then amplified for 15 cycles of PCR using longAmp Taq (NEB M0287S). During amplification PrepX RNAseq index barcode primers were added for each library to enable multiplexing. The amplified library was then cleaned up using PrepX PCR cleanup-8 protocol with magnetic beads (Aline C-1003). The final cDNA libraries were quantified using a high sensitivity dsDNA kit (Thermo Fisher Q32854) on a Qubit 3.0 Fluorometer (Thermo Fisher Q33216). cDNA content and quality were assessed with D1000 (Agilent 5067-5582) or High sensitivity D1000 tape (Agilent 5067-5584, when cDNA was in low amounts) on an Agilent Tapestation 2200 or 4200. For a step-by-step detailed protocol see **Supplementary File 1**.

### Sequencing cDNA libraries

Libraries were sequenced on an Illumina HiSeq 2500 sequencer. Single end-50bp reads were sequenced on a high-throughput flow cell. Libraries of varying concentrations were normalized to be equimolar, the concentrations of which ranged between 2-10nM per lane. All the samples in a flow cell were multiplexed and later separated on the basis of unique prepX indices to yield at least 10 million reads per library. The reads were demultiplexed and trimmed of adapters using the bcl2fastq2 v2.2 pipeline to yield final fastq data files.

### RNA-seq data processing

The *D. melanogaster* genome assembly and gene annotations were obtained from FlyBase version dmel_r6.36_FB2020_05 (Larkin *et al*. 2021). The reads were aligned with RSEM v1.3.3 (Li and Dewey 2011) and using STAR v2.7.6a as read aligner (Dobin *et al*. 2013) we obtained the gene counts in each library. Because some of the tissue-specific biological samples were sequenced in more than one lane or run, and therefore the reads were split into multiple fastq files, the gene counts belonging to the same biological sample were summed. Gene counts in each dataset were normalized with the variance stabilizing transformation (VST) method implemented in the DESeq2 v1.26.0 (Love *et al*. 2014) R package. Further analyses, such as principal component analysis, hierarchical clustering, and differential expression analysis, were performed in R using the VST-normalized counts.

### Differential Expression (DE) analysis

The differential expression analyses were performed with DESeq2 v1.26.0 (Love *et al*. 2014). On the whole ovary dataset, the contrasts tested were early vs mid, and mid vs late stages. For the tissue-specific datasets, three different comparisons were performed. First, to identify differentially expressed genes independently of the stage, all stages of somatic cells were compared to all stages of germ cells. Second, to identify genes up-regulated in a stage-specific manner within each tissue, we compared the expression level at each stage to the mean expression level of the other two stages. Third, we compared germ cells and somatic cells independently at each stage. Genes with a Benjamini-Hochberg (BH) adjusted p-value lower than 0.01 were selected as differentially expressed in the corresponding contrast.

### Functional analysis

The Gene Ontology (GO) and Kyoto Encyclopedia of Genes and Genomes (KEGG) pathways enrichment analyses were performed on the differentially expressed genes with the enrichGO and enrichKEGG functions of the clusterProfiler package (v3.14.3) for R (Yu *et al*. 2012). The GO terms were obtained using the R package AnnotationDbi (Carlson 2015) with the database org.Dm.eg.db v3.10.0. The GO overrepresentation analysis of biological process (BP) was performed against the gene universe of all *D. melanogaster* annotated genes in org.Dm.eg.db, adjusting the p-values with the Benjamini-Hochberg method (BH), adjusted p-value and q-value cutoff of 0.01, and a minimum of 30 genes per term. For the KEGG enrichment analysis, p-values were adjusted by the BH procedure, and an adjusted p-value cutoff of 0.05 was used.

## Data availability

All the raw data are publicly available at NCBI-Gene Expression Omnibus (GEO) database under the accession code GSE172015. The scripts used to process and analyze the data are available at GitHub repository https://github.com/guillemylla/Ovariole_morphogenesis_RNAseq.

## Acknowledgements

This study was supported by National Institutes of Health NICHD award R01-HD073499-01 to CGE, funds from Harvard University, and the Google Cloud Academic Research Credits Program. We thank the flow cytometry and genomics staff at the Bauer Core Facility of Harvard University Faculty of Arts and Sciences (FAS) for advice on FAS sorting and sequencing, the FAS Research Computing group for advice on initial data analysis, and Extavour lab members for discussion.

