## Supplementary File 1 for "Distinct gene expression dynamics in germ line and somatic tissue during ovariole morphogenesis in *Drosophila melanogaster*"

These Supplementary Materials contain the following:

- Detailed Protocols
    I. I. Detailed protocol for staging larvae
    II. Detailed protocol for dissection and dissociation of larval ovaries
    III. Detailed RNA extraction protocol
    IV. Detailed library preparation protocol
- Key Resources Table
- Supplementary References
- Legends for Supplementary Tables S1 through S9 (this document)
- Supplementary Tables S1 through S9 are provided in files S2 through S10
- Supplementary Figures S1 through S12 with Legends (this document)

### DETAILED PROTOCOLS

#### I. Detailed Protocol to stage larvae

1. Day 1: Collect 100 virgin females on the day before egg collection. Set the cross in a 50 ml media bottle with 50 males and leave at room temperature for 12h (overnight) to mate.
2. Make apple juice plates as follows:
    i. Boil 9g bacterial agar (Becton Dickinson catalog # 214050) in 300ml autoclaved distilled water.
    ii. Separately, dissolve 10g Sucrose in 100ml apple juice.
    iii. Mix the two solutions together while stirring with a magnetic stir bar.
    iv. Pour the media into 60x15mm plates once the temperature has cooled down to approximately 50°C.
    v. Cool plates without lids for two hours and then cover with lids and store inverted at 4°C.
    vi. These plates can be used for up to two weeks.
3. Day 2: Remove apple juice agar plates needed for each hour for up to eight hours and allow them to warm up to room temperature.
4. In a glass vial, place some yeast granules and add tap water to cover the granules. There must be a drop of water more than the yeast granules can soak up. This makes a paste of peanut butter consistency.
5. Using a steel spatula, smear a pea-sized amount of paste onto one end of a plate, for all the plates. Optimize this based on the number of flies such that the paste is neither completely consumed nor remains in excess after an hour-long collection.
6. Transfer the cross in the bottle to a 100 ml collection cage and cover it immediately with an apple juice egg-collection plate containing yeast smear. Fasten the setup with two rubber bands.
7. Incubate with the plate at the bottom for one hour at 25°C. All activity from this point until the point of dissection is done in at 25°C. For the first hourly change, tap the bottom of the cage and quickly replace the old plate with a fresh collection plate.
8. Remove any flies stuck to the yeast patch with forceps, crush and discard them in the freezer.
9. After the final egg plate flip transfer the flies back into a bottle using a funnel. Discard the first collection plate. Incubate all the remaining plates at 25°C.
10. Day 3: Start collecting larvae at the end of the second hour egg collection done the previous day. Collect uniformly sized larvae.
11. Transfer around 50 larvae from the same staged egg collection into a vial. Some yeast paste may be carried to the vial during this collection; try to keep this amount constant.
12. Collect until the final hour of the previous days' collection. Incubate the vials containing larvae at 25°C.

**II. Detailed protocol for dissection and dissociation of larval ovaries**

1. Begin dissections at the same time as that of the third plate from the egg collection day. Early-stage dissections take place three days from egg laying (72h AEL), middle (mid) stage four days from egg laying (96h AEL), and larval pupal stage (late) five days from egg laying (120h AEL).
2. Late stages are the easiest stage to locate and dissect, recognizable once the late third instar larvae have immobilized on the side of the vial and have a thickened cuticle. Use a fine wet paintbrush to dislodge them and place into cold 1x phosphate-buffered saline (PBS).
3. For early and middle stages, scrape the soggy layer of food from the vial using a spatula and spread it on a glass petri dish. Under a dissecting stereomicroscope select uniformly sized larvae, wash them in 1xPBS and place them into a fresh glass dish with 1xPBS.
4. Male larvae are easily identified by the two translucent spots that are the testes located at approximately 75% the length of the larval body from the anterior, and by their relatively smaller body size compared to females. Adjusting the light sources to be closer to the stage at the base of the glass dish allows better visualization of ovaries and testes. Discard the males by using forceps to transfer them onto a paper towel or kimwipe.
5. Using forceps, decapitate the larva. Gently squeeze out the inner contents from posterior to anterior using forceps.
6. Pull the larval body away gently with forceps from the fat-body/gut, while holding the body with another forceps. If done properly (after some experience) the larval body and the gut material separate from the fat body lobes. Well-fed and later stages of larvae are easier to dissect than younger stages.
7. The ovaries are located in the middle of the larval fat bodies. Ovaries are in a flower-like circular patch in the center of the fat body. They are around 100-500 μm in size depending on the stage, and appear as transparent tiny dots (early) to large dots (pupal) within the fat body.
8. Using two insulin needles, hold the fat body close to the ovary with one needle and use the other needle to cut closely around the ovary until it is released. Some areas of the fat body may remain initially, which will be removed later when the dissected ovaries are treated with trypsin.
9. Record the number of ovaries acquired in each batch.
10. Start to thaw an aliquot of liberase (stored at -20°C) to room temperature.
11. Dissect approximately 20 ovaries in this way, then place the glass dish on ice. Use a fresh dish for the next round of dissections.
12. Use a 12-well glass plate and add 200μl of trypsin to one well and the contents of the thawed liberase aliquot (~150μl) to the next well.
13. Clear any debris that surrounds the collected ovaries into a separate glass dish using a needle. Using a 2μl pipette filter tip pre-rinsed in trypsin solution, transfer the dissected ovaries into the trypsin well.
14. After transferring the last batch of ovaries into trypsin solution, incubate for 10 minutes in trypsin solution. Meanwhile label two RNA lobind tubes (Eppendorf 8077-230), with a black waterproof marker indicating the genotype of the dissection, the date and the number of ovaries. Add the desired amount (100-

200µl) of Trizol (based on the anticipated number of cells after sorting) to one of the two tubes (tube #1) – this tube will be used to collect cells following FACS (step 18). Leave the second tube (tube #2) empty – this will be used to collect the dissociated cells following enzymatic treatment (step 17). Place both tubes on ice.

15. Using a 2µl filter tip pre-rinsed in liberase solution, transfer the ovaries from trypsin into liberase. Dissociate them using needles until no clusters of cells can be seen. Incubate them in liberase solution for ten minutes, starting the clock when the ovaries are transferred from trypsin (includes time needed to dissociate using needles).
16. Place 200ml of liquid nitrogen in a liquid nitrogen container.
17. Using a 200µl pipette filter tip pre-rinsed in 1xPBS, pipette the tissue in liberase up and down gently ten times to dissociate and resuspend the cells. Transfer the contents to tube #2 and place it on a microtube vortexer for one minute. Meanwhile rinse the well that contained liberase with 1.4 ml of 1xPBS and pipette up and down 10 times. Add this volume into the tube and vortex for an additional 10 minutes.
18. Place the tube on ice, remove one glove (so you can safely touch door handles) and carry the tube on ice, and your liquid nitrogen container, to your FACS facility/machine.
19. Collect cells following FACS into the Trizol in tube #1.

**III. Detailed RNA extraction protocol**

1. Because RNA extraction from FACS-sorted samples involves precious samples, take care to work in an RNase-free environment. Wear lab coat, gloves and safety goggles while working with Trizol and handling RNA samples.
2. Clean the table, ice bucket, pipettes, centrifuge, pellet pestle motor, table top vortexer and tabletop mini-vortex first with 70% ethanol and then with RNase zap (Thermo Fisher AM9780) on tissue paper.
3. Pellet pestles are cleaned first in 100% ethanol and then in nuclease free water. Prior to use the pestles should be sterilized in a glass beaker covered in aluminum foil by autoclaving in a liquid cycle for 30 min.
4. Add 500 µl Trizol into a lo-bind tube, which will be used to pre-rinse the pellet pestle before crushing each sample.
5. Thaw Trizol cell samples (retrieved in tube #1 at step 19 in Protocol II) at room temperature (RT) and place them on ice. Spin in tabletop mini-vortex for 10 seconds.
6. Crush each cell sample with a separate pellet pestle pre-rinsed in the Trizol set aside for this purpose in step 4.
7. Crush cells in Trizol with a pre-rinsed pellet pestle. Add an equal volume of 100% ethanol, mix with pestle and vortex briefly. Spin samples briefly in tabletop mini-vortex.
8. Pipette the sample onto a zymo spin column.
9. Wash column in 400 µl RNA wash buffer. Centrifuge at 10,000 g for 1 minute at RT.
10. Thaw DNase (6unit/µl) from storage at -20°C. Mix 5 µl DNase with 35 µl DNase digestion buffer per sample. Add 40 µl of DNase mix to each column.
11. Incubate at RT for 15 minutes. Centrifuge at 10,000 g for 1 minute at RT..
12. Add 400 µl RNA pre-wash buffer to the column and centrifuge. Repeat this step one more time.
13. Add 700 µl of RNA wash buffer. Centrifuge twice to completely remove the buffer from the column, each time at 10,000 g for 1 minute at RT..
14. The column-bound RNA is eluted in two steps using nuclease-free water. Add 25 µl water and incubate for 15 minutes at RT. Centrifuge at 10,000 g for 1 minute at RT. Add another 20 µl of water to elute a second time.
15. The library preparation protocol below (**IV**) requires RNA in a 50 µl volume; an additional 5 µl is reserved for quality control measurements.
16. Quantify RNA first in Nanodrop RNA-40 measurement with 1.5 µl of the sample. Then use Qubit high sensitivity RNA kit to quantify 1µl of the sample (10µl Std1/2+190 Buffer/Reagent, 1 µl sample+199 Buffer/Reagent).
17. Use a high-sensitivity RNA tape to quantify and measure the RNA integrity (RIN) in a Tapestation. Use the standard protocol for high-sensitivity RNA quantification using an electronic ladder for estimation of size.

### IV. Detailed library preparation protocol (Takara Apollo system)

This is the Wafergen/Takara protocol. Low-throughput protocols process a maximum of eight samples at a time in the Apollo liquid handling unit. Label the individual tubes in the strip with the respective sample identifiers and make sure the marked wells are in the same 1-8 sequence direction at all times. Use the protocol images to ensure accurate placement of strip tubes and double check tube placements regularly. RNA extraction, Poly A selection and Library preparation should be done on the same day.

#### *IV.i. Poly A selection protocol*

1. Verify that the volumes of RNA samples are at least 50 µl using micropipettes and pipette them into a strip tube (Fisher scientific 14-222-252). Label tube as "RNA samples". Spin down in a tabletop mini-centrifuge and keep on ice until step 7.
2. Wipe the inner surface of the Apollo system with RNase zap (Thermo Fisher AM9780) and 70% Ethanol.
3. Ensure the trash container is empty. The machine shows an error when trash is full.
4. Cool the Apollo machine to 4°C using the standard protocol 'cooling' function.
5. Place empty reservoirs (Apollo 640087) in Block-6 row 1 and 2.
6. Fill Block-5 row 1,2 and 3 with filter tips (Apollo 640084).
7. Place a new microplate in Block-2.
8. In two new reservoirs, add 10 ml (protocol says 4/5ml) of Reagent 1 and place it in row 1 and 10ml of reagent 2 in row 2.
9. Place empty 8 strip tubes in Block-3 row 4 and 5 and in Block-4 row 1, 2 and 3.
10. Label an empty strip tube as "products" and place it in Block-3 row 8. This strip tube will contain the poly-A selected mRNA at the end of this run. Label each tube with the sample names.
11. Aliquot 80µl of Reagent 3 (it is a clear solution) into an 8 well strip tube into wells corresponding to the RNA sample. Label as "Reagent 3" to distinguish it from RNA samples.
12. Place Reagent 3 strip tube into Block-3 row 3.
13. Gently pipette the magnetic bead reagent 4 until it is uniformly resuspended. Aliquot 15 µl into the same number and locations of the strip tube containing RNA sample.
14. Place the RNA sample tube in Block-3 row 1.
15. At last place the magnetic bead reagent 4 in Block-3 row 2. Place retainers for Blocks 3 and 4 and lock them.
16. Restart the machine by switching off and on. On the touchscreen navigate to the latest version of PolyA8 protocol ("User Maintenance" > "PrepX_PolyA8_betaV2").
17. The run lasts for 45 minutes, during which time the reagent mixes for the subsequent library preparation protocol (**IV.ii**) should be set up.
18. Remove strip tubes while checking for uniform volumes. Note any discrepancies.

19. Check the volume of the “product” tube, cap it and spin it in a minispin. The volume should be approximately 19µl. Cap and store on ice until the next step. This mRNA “product” strip tube will be used as “sample” in the cDNA library preparation step.

#### *IV.ii. PrepX mRNA8 Library preparation protocol.*

1. Prepare the RNase III mix and Reverse Transcription (RT) reaction mix for the number of mRNA samples and an additional sample.
2. In a RNase free tube mix 2µl each of RNase Buffer III and RNase III enzyme (Thermo Fisher 18080093) per sample required. Pipette gently and give a brief spin on a tabletop mini-centrifuge. Leave the tube on ice.
3. Mix the following reagents per reaction to make the RT reaction mix and place it on ice:
    - 5X First strand buffer 16µl
    - 0.1M DTT 08µl
    - dNTP 04µl
    - Superscript III Reverse Transcriptase 02µl
    - Murine RNase inhibitors 01µl

Setting the apollo system blocks:

4. Place empty strip tubes in Block-3 rows 1 and 2 and in Block-4 row 6.
5. Label a strip tube as products and place it in Block-3 row 5.
6. Fill filter tips in Block-5 rows 1-7, fill black piercing tips (Apollo 640085) in row 12.
7. Fill 1.1 ml tube strips in Block-1 row 1,2 and 3.
8. Place a used microplate in Block-2.
9. Cool the Apollo machine to 4°C using standard protocol ‘cooling’ function.
10. Place empty reservoirs in position 2 and 4 of Block-6. Add 15 ml of 100% molecular grade ethanol (Sigma E7023) in reservoir 3 and 15 ml of nuclease free water (part of Takara library prep kit 640096) in reservoir 1.
11. In a new strip tube, aliquot 4 µl of RNase III mix into each tube corresponding to the sample and place it in Block-4 row 5.
12. In a new strip tube, aliquot 31 µl of RT reaction mix into each corresponding sample tube and place it in Block-4 row 7.
13. Gently pipette A-line beads and aliquot 200µl into a fresh strip tube at the corresponding wells to that of the sample and place it in Block-4 row 8.
14. Thaw blue enzyme and orange adapter/primer strips of the PrepX mRNA8 kit (Takara 640096) on ice. The number of these strips is the same as the sample number. Flick the bottom of tubes to dislodge liquid and mix uniformly. Spin down and place it back on ice.
15. Sometimes a solid precipitate might be visible in the enzyme strip tubes. It generally dissolves after flicking. Do not use the strip if it is not soluble after thawing and mixing.
16. Place the Blue strip tubes with the arrow pointing up in Block4 rows 9-12 in columns corresponding to the mRNA samples.

17. Place the orange strip tubes in Block-4 from row 1-4 in columns corresponding to the mRNA sample tubes.
18. Verify the filter and piercing tips in Block-5, 1.1ml tubes in Block-1, mock microplate in Block-2, Reservoir 3 with 100% ethanol and Reservoir-1 with Nuclease free water, Reservoirs 4-5 empty place holders, three sets of empty tubes (one for cDNA products) and mRNA sample tube in Block-3 and in Block-4 Blue and orange strips, one empty strip tube, RT, RNase III and bead strip tubes in specified locations.
19. Lock Block-3 and 4 with retainer plates.
20. In the touchscreen control select User maintenance > Run the protocol ‘PrepX_mRNA8_200bp_BetaV1.scb. The screen does not show any progress bar. This program runs for 5 hours.
21. cDNA after this step is fairly stable and can be processed the next day. Keep at 4°C until processing.

#### *IV.iii. PCR Amplification of the Libraries.*

1. Check the volume of the “product” tube, cap it and spin it in a minispin. The volume should be around 19µl. Cap and store on ice until the next step.
2. Prepare PCR master mix with 25µl long Amp Taq (NEB M0323S) and 2.5 µl of SR primer per sample on ice.
3. Add unique index primers (PrepX RNAseq index 1-48) to each of the cDNA products and add the PCR master mix. Make up the total volume in each tube to 50µl with Nuclease free water.
4. Mix well and spin down. Place the strip tube in a PCR machine and run a 15 cycle PCR amplification reaction as shown below:

| **Temp** | 94℃ | 94℃ | 60℃ | 65℃ | 65℃ | 10℃ |
|---|---|---|---|---|---|---|
| **Time** | 60 sec | 30 sec | 30 sec | 30 sec | 7 min | hold |
| Cycles |  | -------------------- 14 x ------------------- |  |  | extension |  |

6. After completion of PCR cool the samples and spin down.
7. PCR Cleanup using the Apollo system PrepX_PCR clean up8 protocol.
8. Cool the Apollo machine to 4°C using standard protocol ‘cooling’ function.
9. Place filter tips in Block-5 row 1. Block-6 is similar to the library preparation protocol, empty reservoirs in position 2 and 4 of Block-6. 10 ml of 100% molecular grade ethanol (Sigma E7023) in reservoir 3 and 15 ml of nuclease free water (part of Takara library prep kit 640096)) in reservoir 1.
10. Place empty strip tubes in Block-3 row 2 and 3. Label a strip tube as “cleaned cDNA product” and place it in row 4. Place the amplified cDNA samples in row 1.
11. Aliquot 50 µl of A-line beads into a fresh strip tube at corresponding locations to the samples. Place this tube in the end of the set up.
12. Place retainers on Block-3 and 4 and lock it.

13. Run Utility apps> PCR cleanup 8 protocol on touch screen. The run lasts 20 minutes.
14. Check volumes of the cleaned ‘product’ tube. Spin down and place on ice.
15. Run Qubit to quantify the cDNA with the high sensitivity DNA kit. Use a Tapestation to run the gel and measure cDNA quantity using ladder and high sensitivity tape (Agilent 5067-5579). Depending on the qubit quantification, higher concentrations use DNA 1000 tape (Agilent 5067-5582).
16. Transfer the cDNA in strip tubes to a lobind RNA tube (Eppendorf 8077-230) and label them with details about the sample-genotype, date of cDNA prep, it is a cDNA library, the concentration of the sample, and the volume of sample remaining after quantification. Store at -80°C until it is ready to be sequenced.
17. Calculate the total molar concentration of the lane and the dilution needed for each library to make it equimolar. Mix the volumes in a single lobind tube and submit to the sequencing facility.

### KEY RESOURCES TABLE

| REAGENT or RESOURCE | SOURCE | IDENTIFIER |
|---|---|---|
| **Chemicals** | | |
| Hoechst 33342 | Thermo Fisher | Cat# H1399 |
| Dulbecco’s Phosphate Buffered Saline PBS | Thermo Fisher | Cat# 14190144 |
| Trypsin 0.25% | Thermo Fisher | Cat# 25200056 |
| Liberase 2.5% | Sigma | Cat# 5401119001 |
| Trizol | Thermo Fisher | Cat# 15596206 |
| Ethanol molecular 200 grade | Sigma | Cat# E7023 |
| Nuclease free water | Thermo Fisher | Cat# 10977015 |
| Magnetic Beads | A-line | Cat# C1003 |
| Triton X100 | VWR | Cat# 97062-208 |
| **Critical Commercial Assays** | | |

| | | |
|---|---|---|
| Zymo RNA Micro-prep kit | Zymo Research | Cat# R2060 |
| Superscript III Reverse Transcriptase | Thermo Fisher | Cat# 18080093 |
| Takara PrepX PolyA-8 | Takara | Cat# 640098 |
| Takara PrepX mRNA-8 | Takara | Cat# 640096 |
| Qubit RNA HS Assay Kit | Thermo Fisher | Cat# Q32852 |
| Qubit DNA HS Assay Kit | Thermo Fisher | Cat# Q32854 |
| LongAmp Taq DNA Polymerase | New England BioLabs | Cat# M0287S |
| **Deposited Data** | | |
| Raw and analyzed data | This paper | GEO: GSE172015 |
| **Experimental Models: Organisms/Strains** | | |
| *D. melanogaster*: bab1 GAL4: *w[*]; P{w[+mW.hs]=GawB}bab1[PGAL4-2]/TM6B, Tb[1]* | Bloomington Drosophila Stock Center | BDSC:6803; FlyBase:ID FBst0006803 |
| *D. melanogaster*: nos GAL4: *P{w[+mC] =UAS-Dcr-2.D}1, w[1118]; P{w[+mC]= GAL4-nos.NGT}40* | Bloomington Drosophila Stock Center | BDSC:25751; FlyBase: ID FBst0025751 |

| | | |
|---|---|---|
| *D. melanogaster*: *w1118, P{UAS Stinger}* | (BAROLO *et al.* 2000) | UAS Green Stinger on X |
| **Instruments** | | |
| MoFlo Astrios EQ Cell sorter | Beckman Coulter | B25982 |
| Motorized pellet pestle | Kimble | Cat# 749540-0000 |
| NanoDrop | Nanodrop | ND1000 |
| Qubit 3.0 Fluorometer | Thermo Fisher | Cat# Q33216 |
| Tapestation | Agilent | 2200/4200 |
| PCR Thermal cycler | Bio-Rad | C1000 |
| Illumina Hi Seq | Illumina | 2500 |
| **Consumables** | | |
| Insulin Syringe | Becton Dickinson | Cat# 328418 |
| RNA lo-bind tubes | Eppendorf | Cat# 8077-230 |
| High Sensitivity RNA ScreenTape | Agilent | Cat# 5067-5579 |

| | | |
|---|---|---|
| High Sensitivity DNA ScreenTape | Agilent | Cat# 5067-5584 |
| DNA 1000 ScreenTape | Agilent | Cat# 5067-5582 |
| Axygen PCR 8-strip tubes | Fisher Scientific | Cat# 14-222-252 |
| Apollo Filter tips 300027 | Takara | Cat# 640084 |
| Apollo Piercing tips 300028 | Takara | Cat# 640085 |
| Apollo Reservoirs 300031 | Takara | Cat# 640087 |
| **Software and Algorithms** | | |
| Drosophila melanogaster genome version | (Larkin *et al.* 2021) | Dmel_r6.36_FB2020_05 |
| RSEM v 1.3.3 | (Li and Dewey 2011) | https://deweylab.github.io/RSEM/ |
| STAR aligner v 2.7.6a | (Dobin *et al.* 2013) | https://github.com/alexdobin/STAR |
| DESeq2 v 1.26.0 | (Love *et al.* 2014) | https://bioconductor.org/packages/release/bioc/html/DESeq2.html |
| Bcl2fastq2 v2.20 | | https://support.illumina.com/downloads/bcl2fastq-conversion-software-v2-20.html |

### SUPPLEMENTARY TABLE LEGENDS

**Supplementary Table S1**: RNA-seq sample metadata, including the sample name, biological and technical replicate information, tissue, stage, cDNA concentration, PrepX Index used, number of ovaries used, number of cells counted by FACS (not applicable to whole ovary samples), number of raw reads, aligned reads, and percentage of aligned reads as reported by the RSEM summary.

**Supplementary Table S2**: Differentially expressed genes (padj<0.01) between the consecutive developmental stages of whole ovary libraries. The column “Contrast” indicates whether the gene was found differentially expressed in early vs mid stages or mid vs late stages, and the column “Up_in” indicates which library the gene was overexpressed in.

**Supplementary Table S3**: Differentially expressed genes (padj<0.01) at each stage compared to the other two stages of the whole ovary. The column “Stage_up” indicates which stage the given gene was overexpressed in.

**Supplementary Table S4**: Differentially expressed genes (padj<0.01) between germ cells and somatic cells at all studied stages. The column “Up_in” indicates whether the gene was found upregulated in germ cells or somatic cells.

**Supplementary Table S5**: Differentially expressed genes (padj<0.01) between germ cells and somatic cells at each individual stage. The column “Up_in” indicates whether the gene was found upregulated in germ cells or somatic cells, and the column “Stage” indicates the stage (early, mid or late) in which the test was performed.

**Supplementary Table S6**: Differentially expressed genes (padj<0.01) between the consecutive developmental stages of somatic cell libraries. The column “Transition” indicates whether the gene was found differentially expressed in the transition from early to mid-stage or from mid to late stage, and the column “Up_Down” indicates whether the gene was up or down regulated in the given transition.

**Supplementary Table S7**: Differentially expressed genes (padj<0.01) at each stage compared to the other two stages of the somatic tissue library. The column “Stage_up” indicates the stage that a given gene was found overexpressed at.

**Supplementary Table S8**: Differentially expressed genes (padj<0.01) between the consecutive stages of germ cell libraries. The column “Transition” indicates whether the gene was found differentially expressed in the transition from early to mid-stage or from mid to late stage, and the column “Up_Down” indicates whether the gene is up or down regulated in the given transition.

**Supplementary Table S9**: Differentially expressed genes (padj<0.01) at each stage compared to the other two stages of the germ cell library. The column “Stage_up” indicates the stage at which the given gene was found to be overexpressed.

### SUPPLEMENTARY FIGURES AND LEGENDS

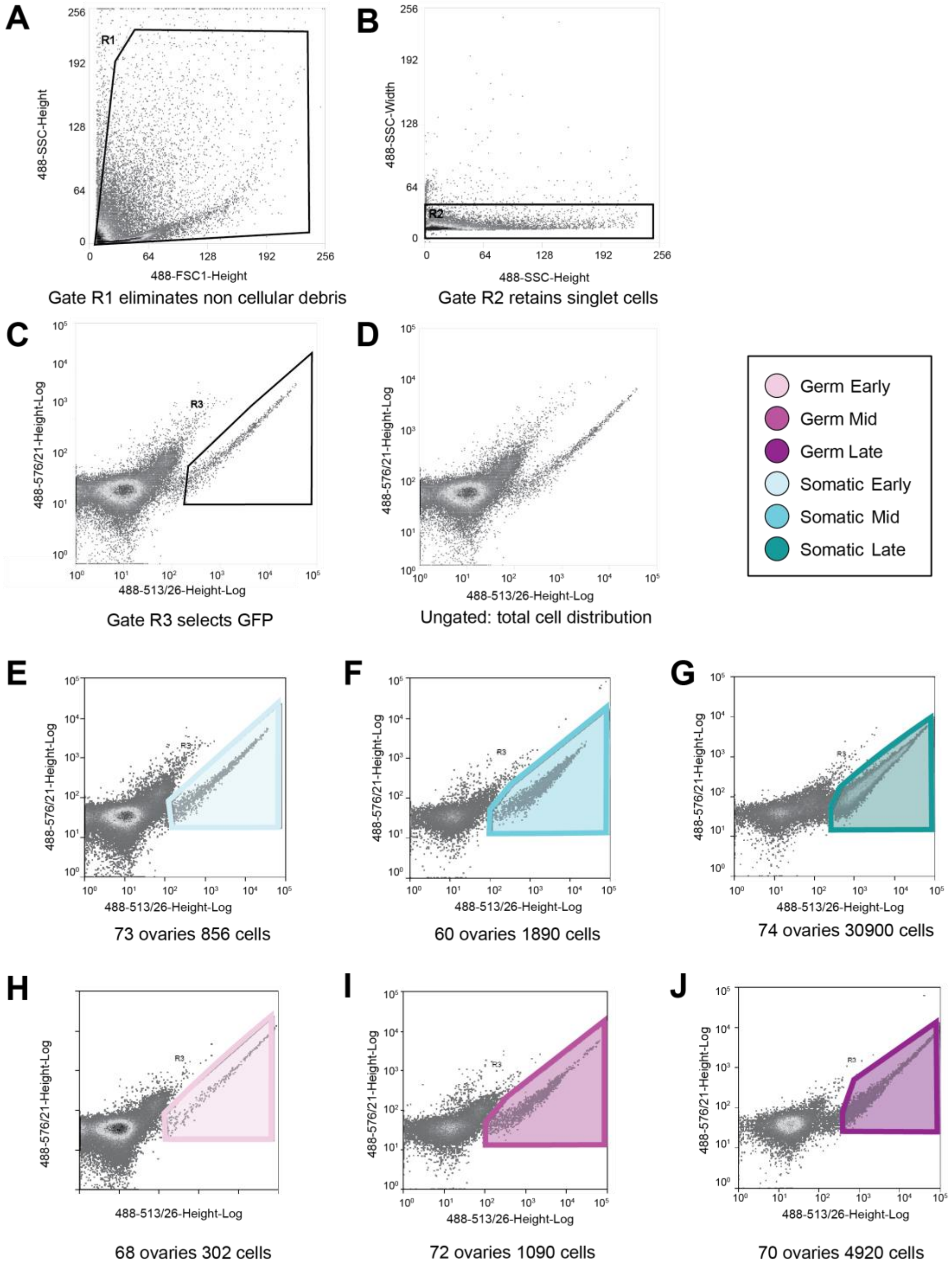


**Supplementary Figure S1.** Number of aligned reads in each of three biological replicates of whole ovary RNA-seq samples. The Black dashed line: five million reads; red dashed line: ten million reads.

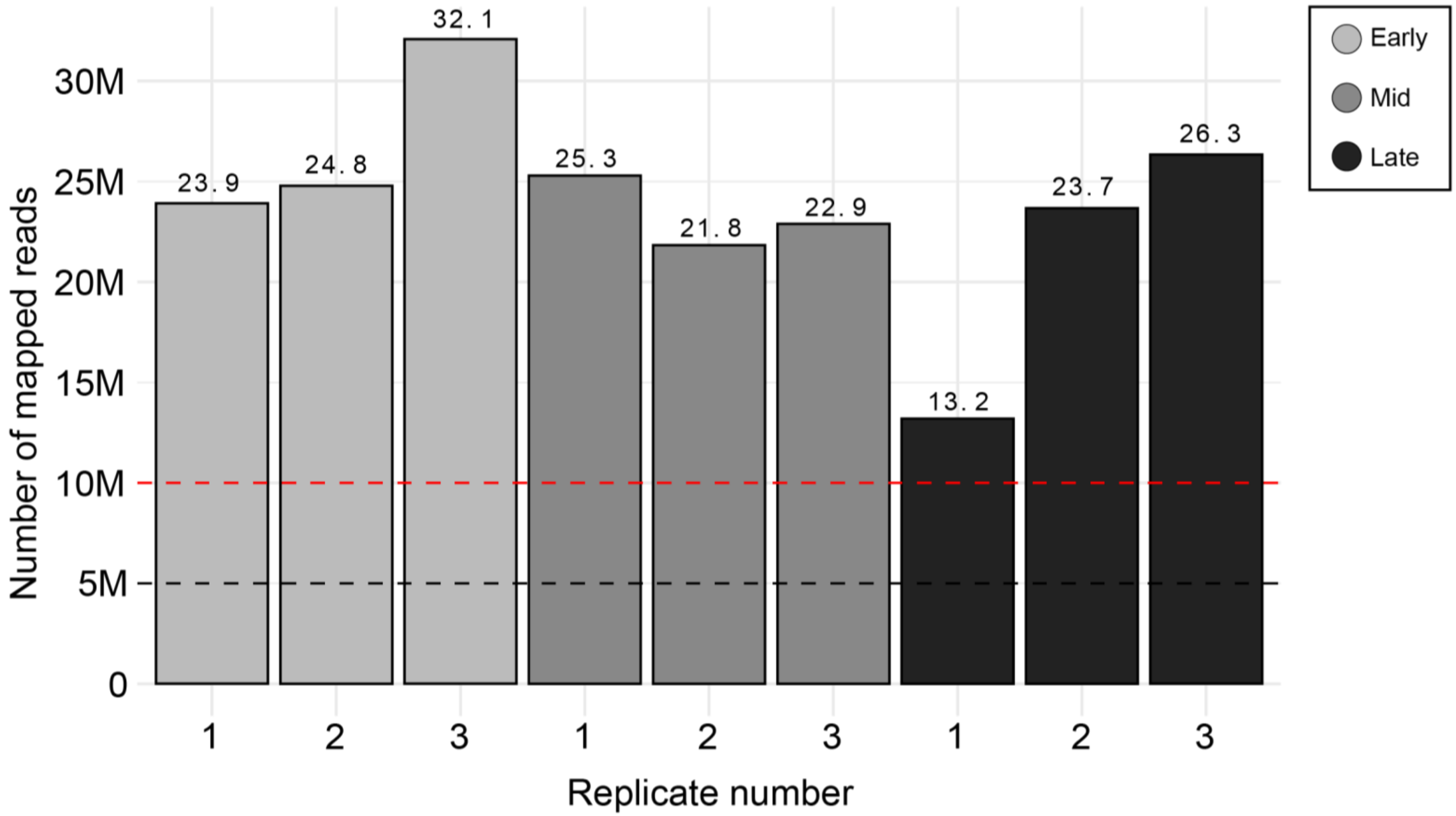


**Supplementary Figure S2.** FACS cell density plots resulting from of sorting GFP-positive cells from dissociated ovaries in a representative plot. **A)** Elimination of cellular debris using R1 (cells out of R1) gate. **B)** Elimination of non-singlets (cells out of R2) using R2 gate. **C)** Selection of GFP-positive cells through R3 (cells inside R3) gate**. D)** Ungated plot showing distribution of all cells. **E-J)** Representative R3 gated plots showing number of GFP-positive cells for similar number of ovaries.

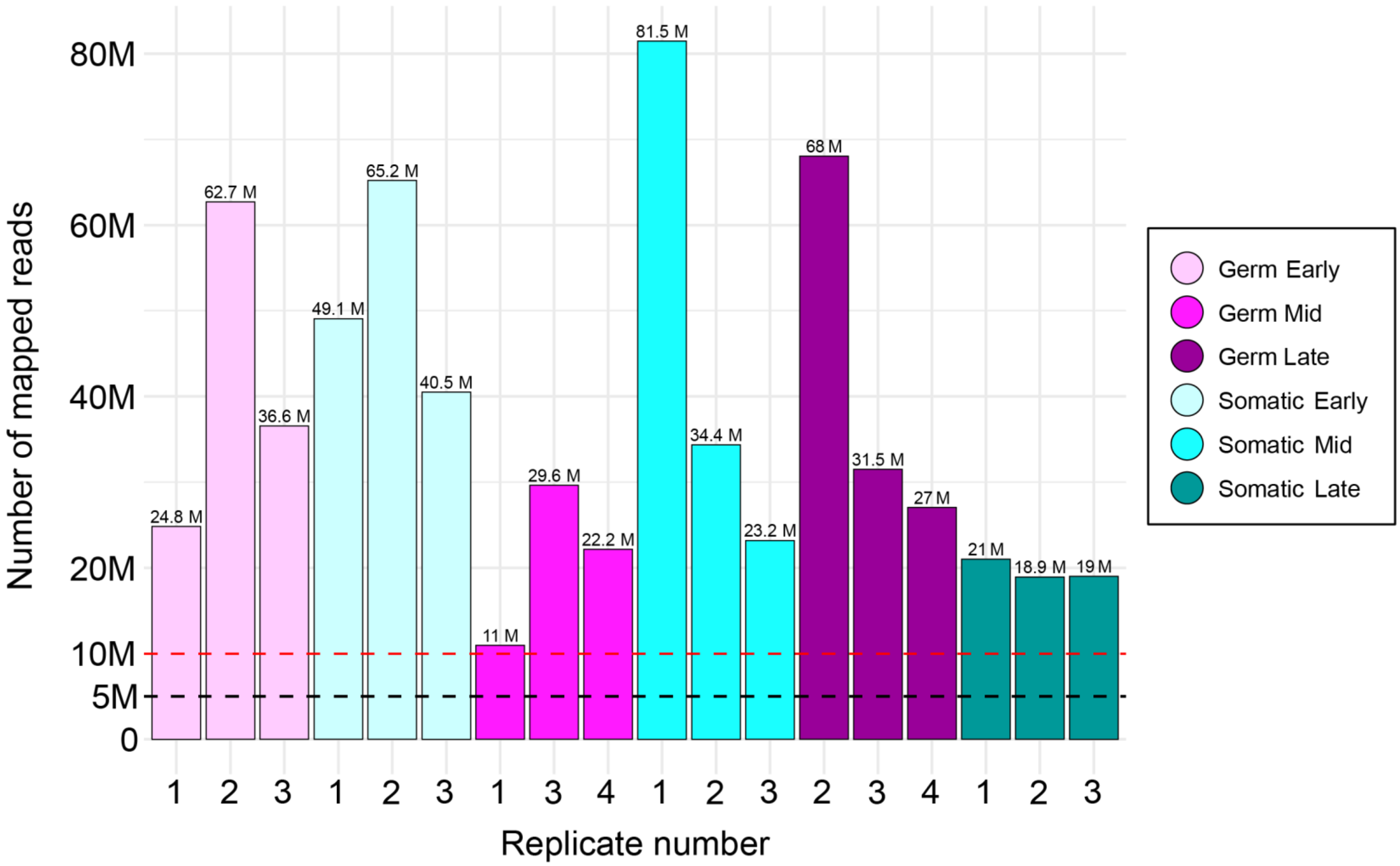


**Supplementary Figure S3**. Number of aligned reads in each of three tissue-specific biological replicate samples used for the analyses presented in this study. Black dashed line: five million reads; red dashed line: ten10 million reads.

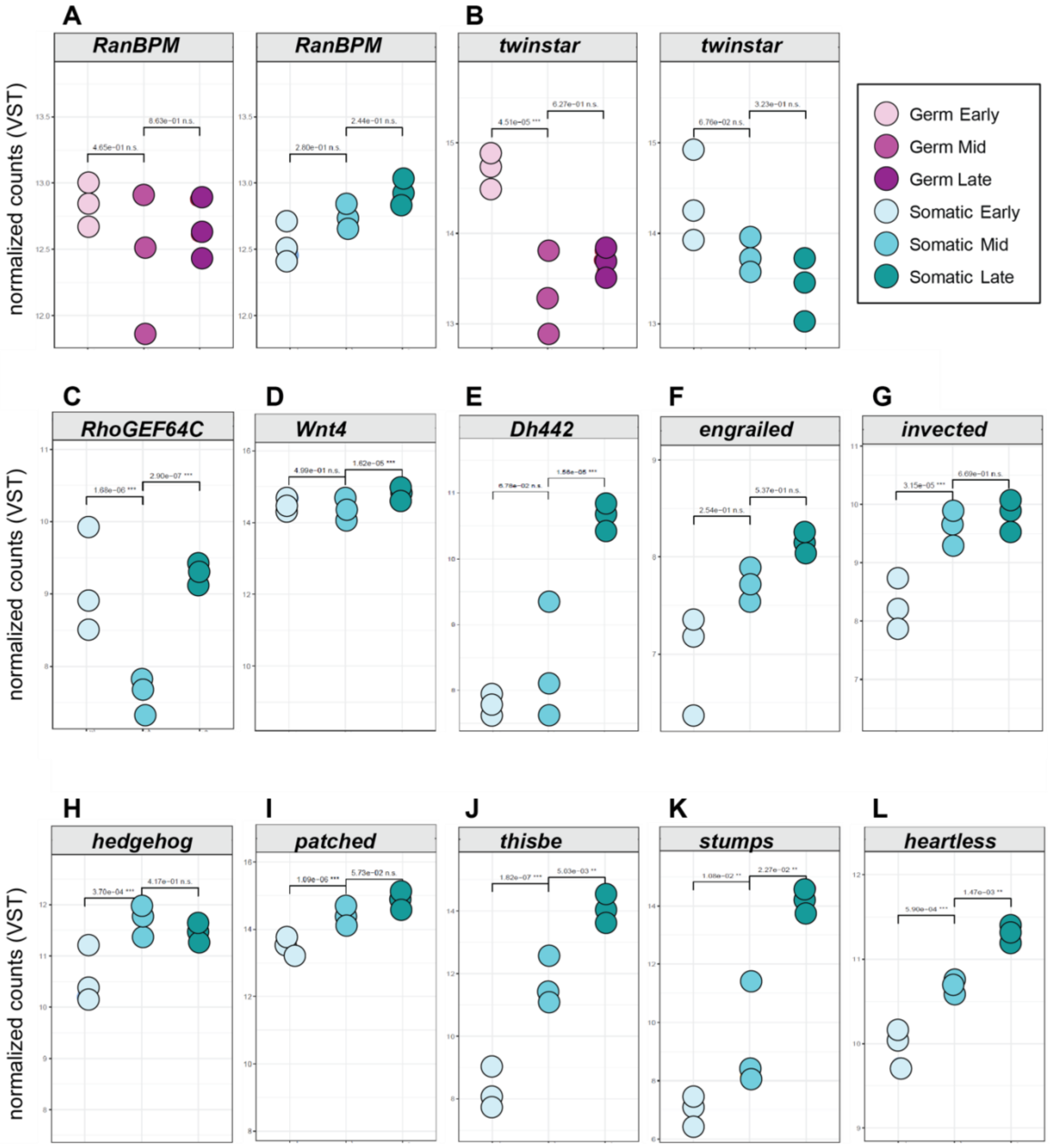


**Supplementary Figure S4**. Dot plots of genes differentially expressed across stages. Expression in VST counts of genes in each tissue-specific RNA-seq library. . The adjusted p-values shown were calculated in the differential expression analysis with DESeq2. *p-value<0.05, **p-value<0.01, ***p-value<0.001, n.s. not significant

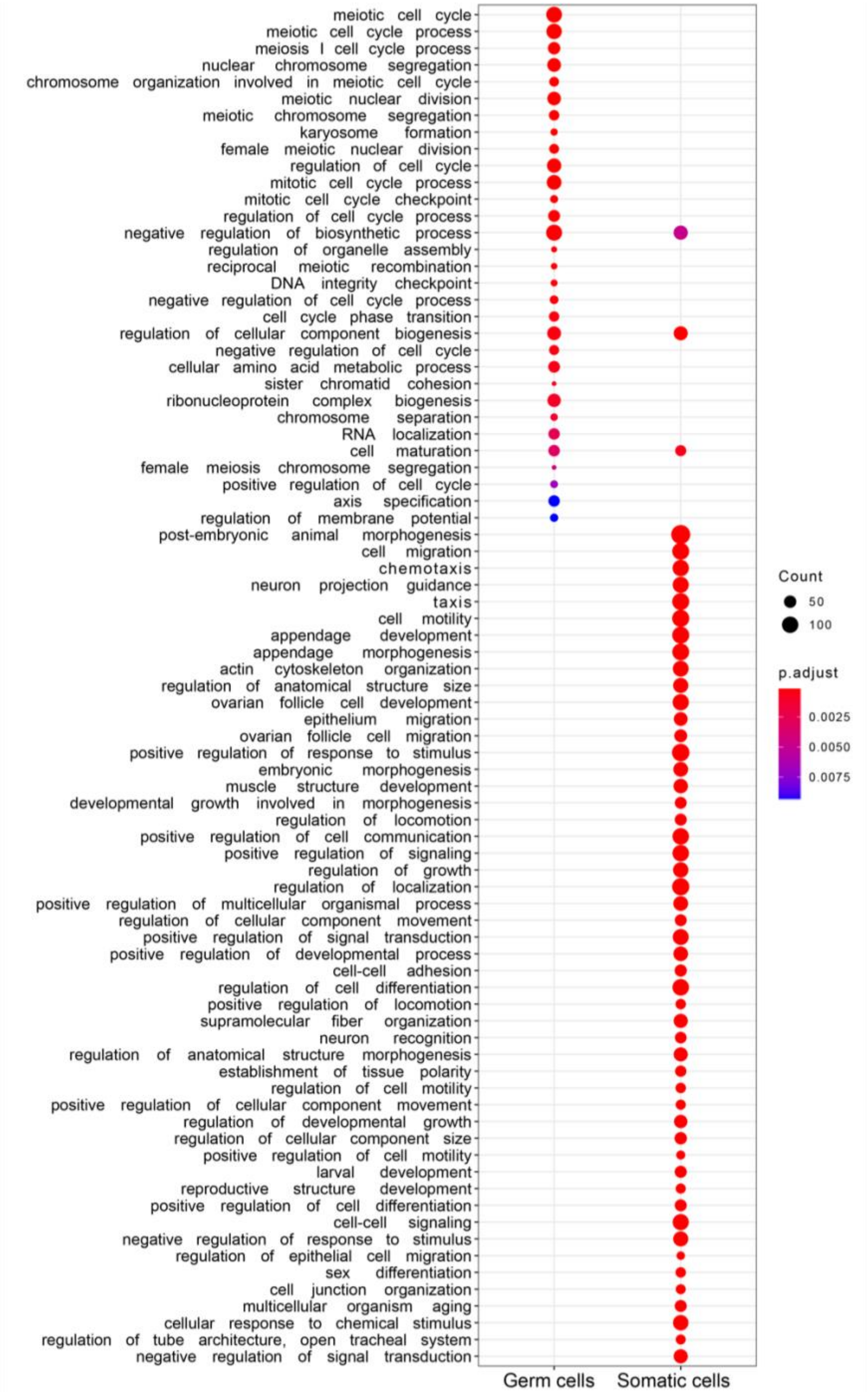


**Supplementary Figure S5.** Gene Ontology terms (GO-terms) for Biological Process level 4 that were significantly enriched (BH adjusted p-value<0.01, minimum number genes with the term=30) within the set of genes differentially expressed (BH adjusted p-value<0.01) between germ cells and somatic cells. The circle size is proportional to the number of genes with the GO-term in the corresponding gene set. The color of the circle indicates the adjusted p-value.

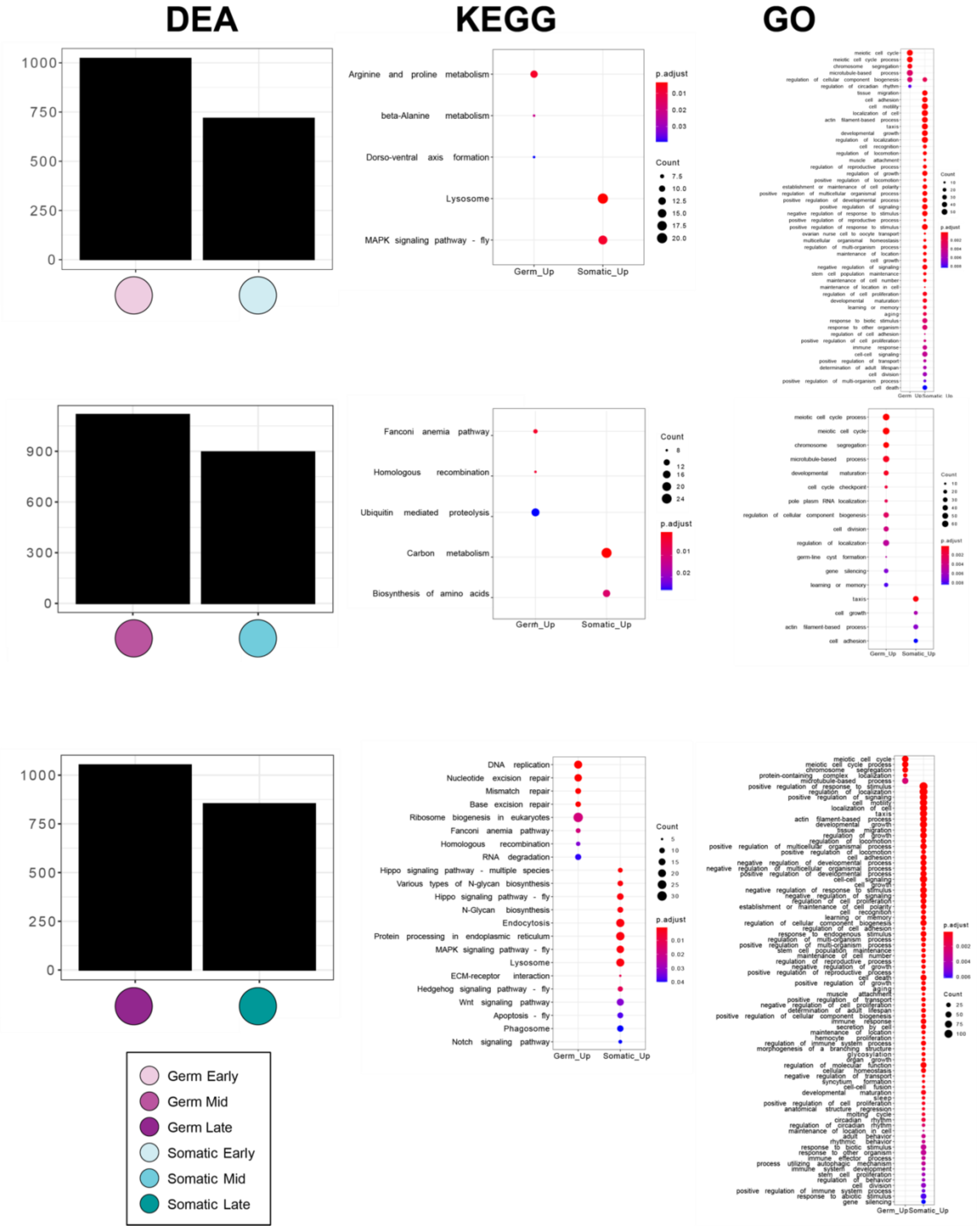


**Supplementary Figure S6.** Number of significantly upregulated genes in germ cells and somatic cells at each of the three stages, and the KEGG pathways and GO-term identified as significantly enriched (BH adjusted p-value<0.05 for KEGG and BH p-value<0.01 for GO) within each set of upregulated genes.

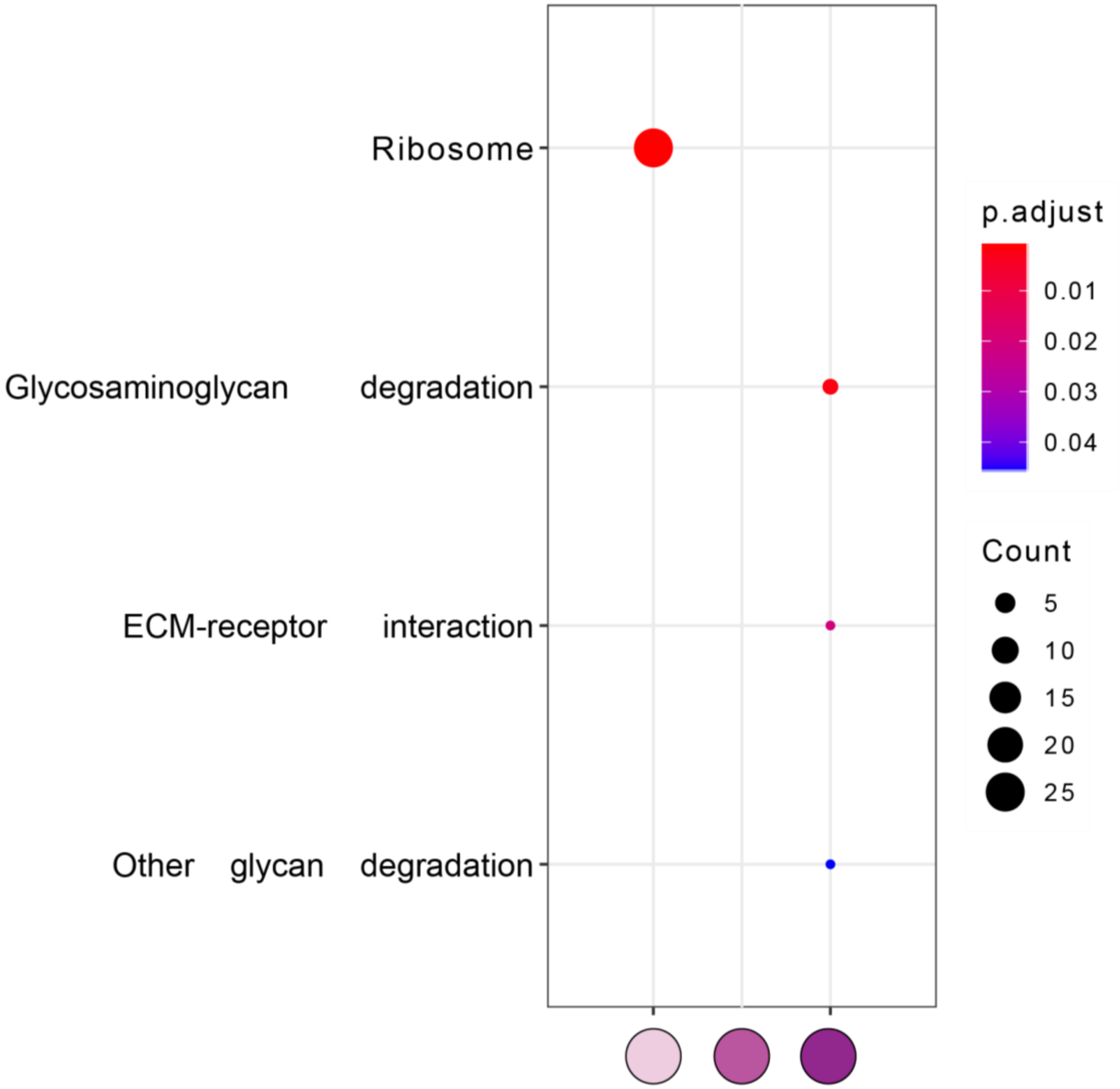


**Supplementary Figure S7.** Significantly enriched (BH adjusted p-value<0.05) KEGG pathways within the sets of genes significantly overexpressed (p-value<0.01) in each stage of the germ cell libraries. The circle size is proportional to the number of genes with the GO-term in the corresponding gene set. The color of the circle indicates the adjusted p-value.

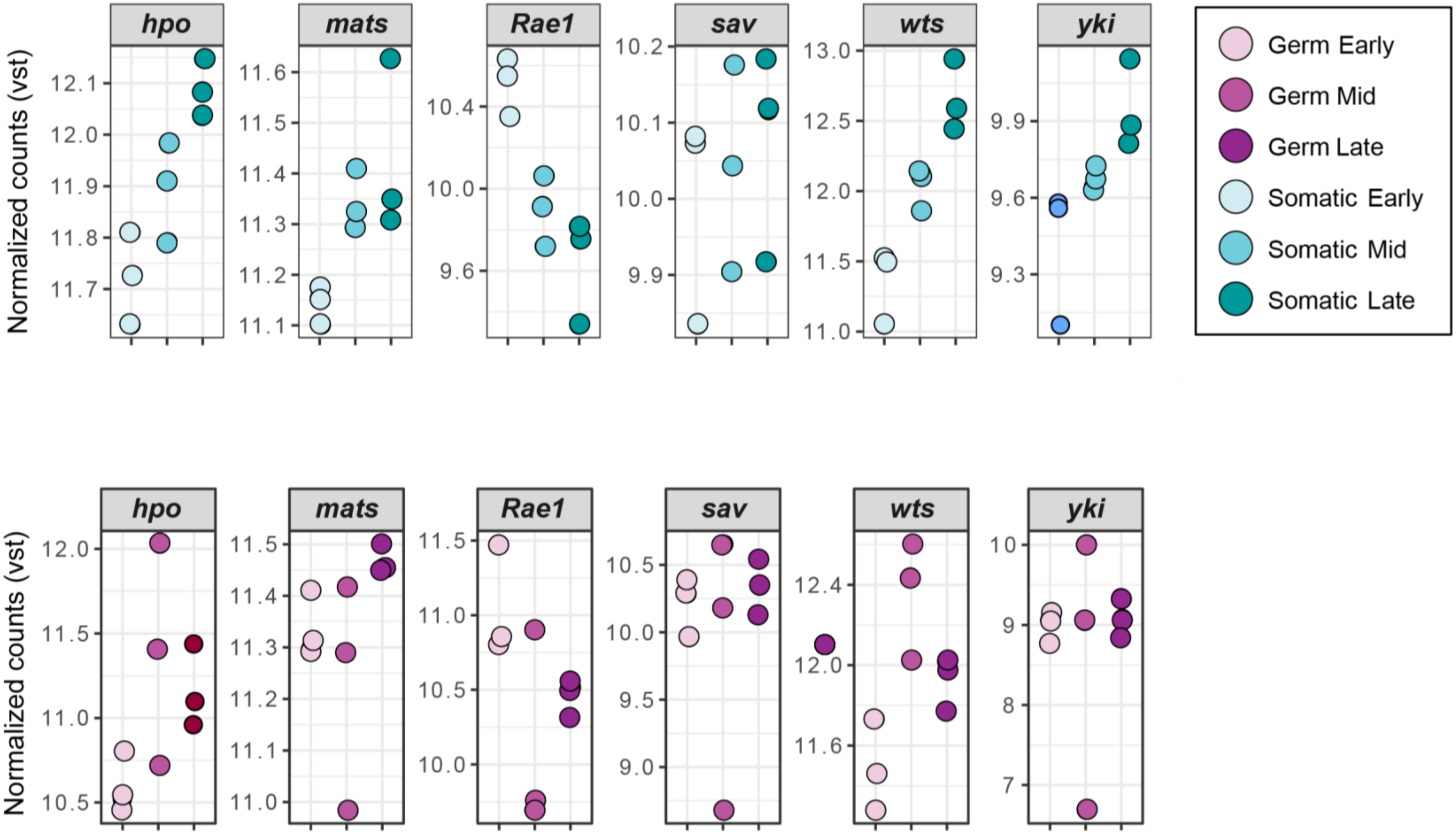


**Supplementary Figure S8**. Expression in VST counts of the Hippo signaling pathway core component genes according to FlyBase (FBgg0000913) in each tissue-specific RNA-seq library. Note that y axes differ slightly between plots

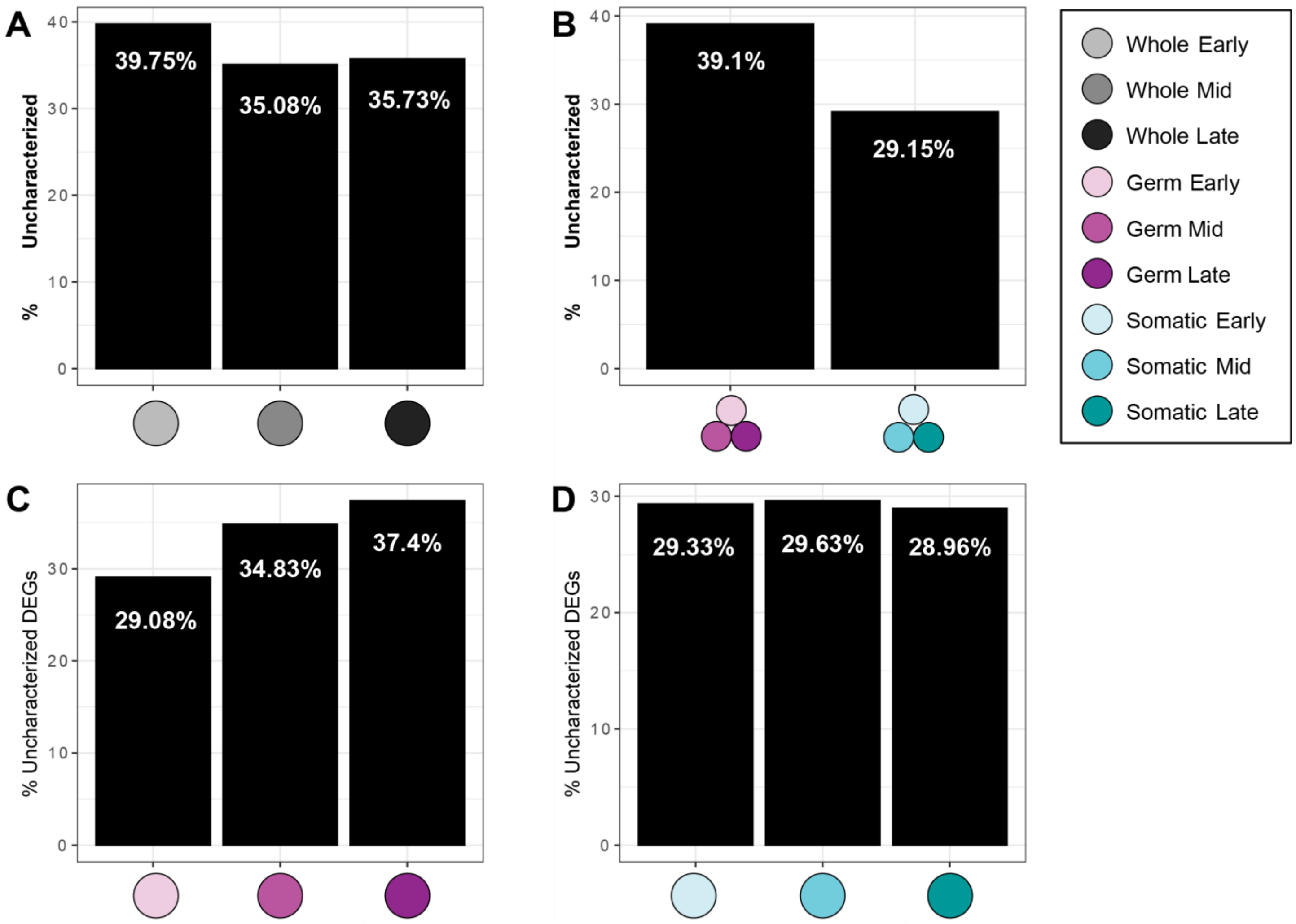


**Supplementary Figure S9.** Uncharacterized genes in differential expression comparisons.

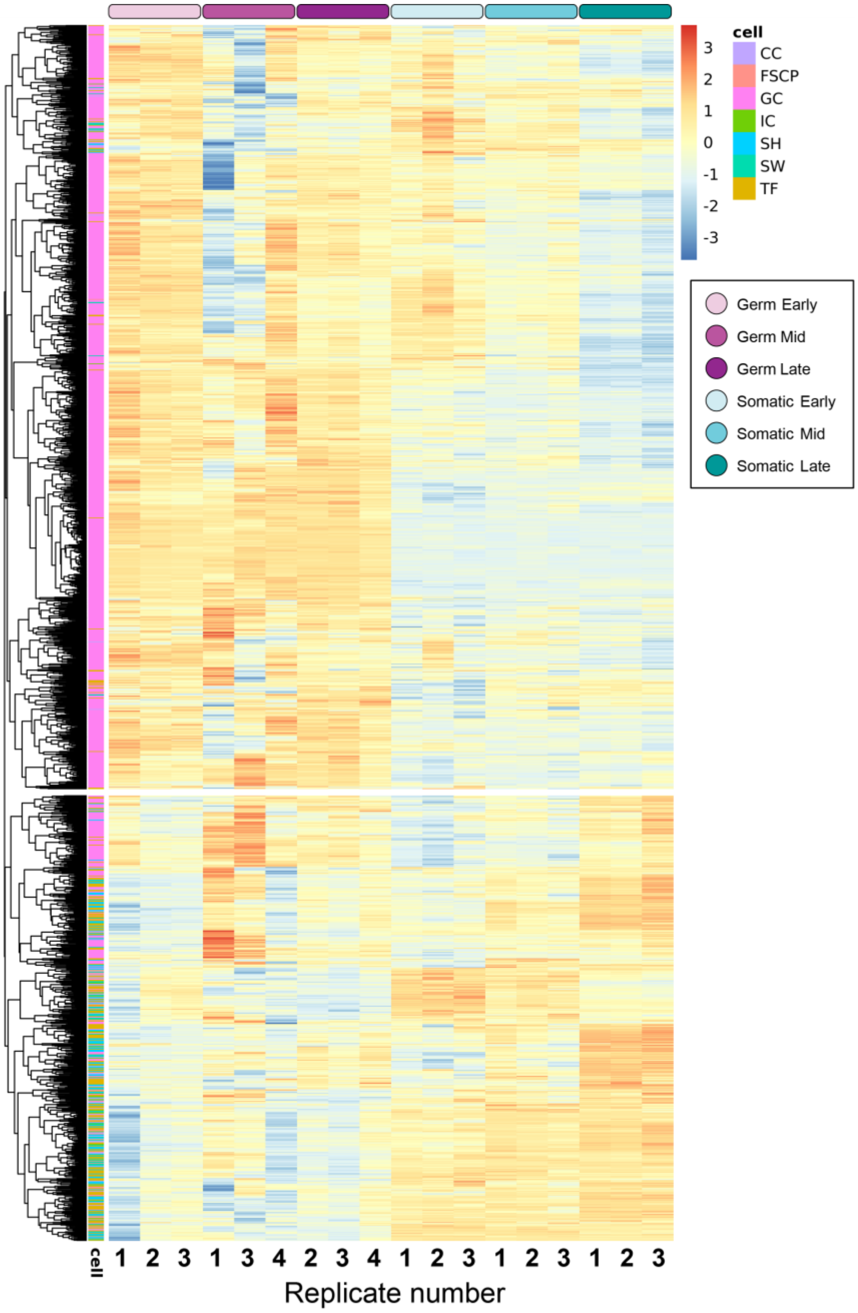


**Supplementary Figure S10.** Expression across our dataset of the cell markers exclusive of a single cell type obtained from Slaidina *et al.* (2020). The color of the first indicates the cell type of which each gene is a marker of (CC: cap cells, FSCP: follicle stem cells, GC: germ cells, IC: intermingled cells, SH: sheath cells, SW: swarm cells, TF: terminal filament). Genes are clustered based on hierarchical clustering and separated into two groups using the cuttree function which resulted in the separation of the germ cell markers from somatic markers. The expression of each gene across samples is represented as a row-wise Z-Score value of the VST-normalized counts from high(red) to low (blue).

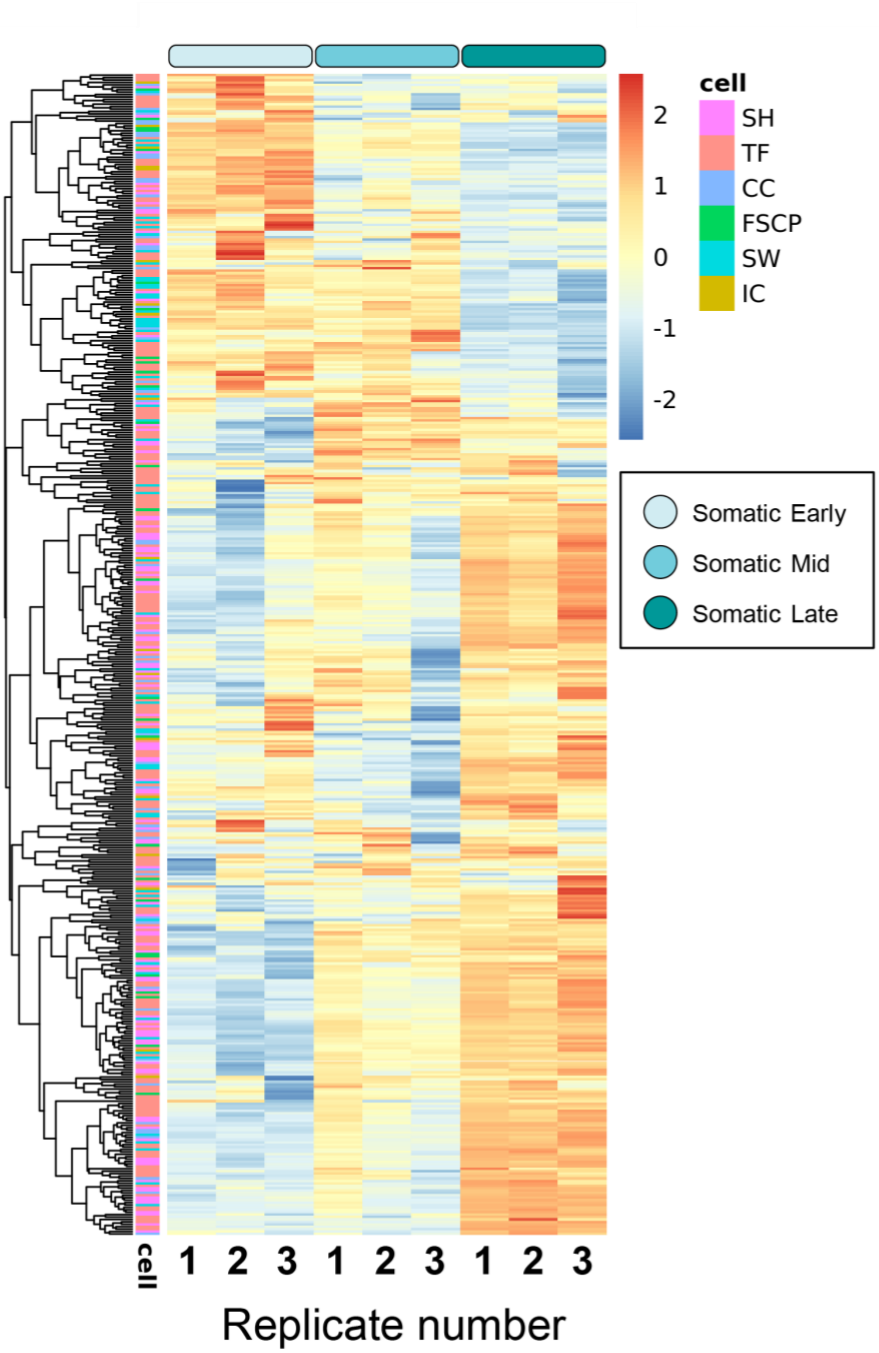


**Supplementary Figure S11.** Expression across stages in somatic cell libraries of the somatic markers exclusive of a single cell type obtained from Slaidina *et al.* (2020). The color of the leftmost column indicates the cell type suggested by each marker gene (CC: cap cells; FSCP: follicle stem cells; GC: germ cells; IC: intermingled cells; SH: sheath cells; SW: swarm cells; TF: terminal filament). Genes are grouped based on hierarchical clustering. The expression of each gene across samples is represented as a row-wise Z-Score value of the VST-normalized counts from high (red) to low (blue).

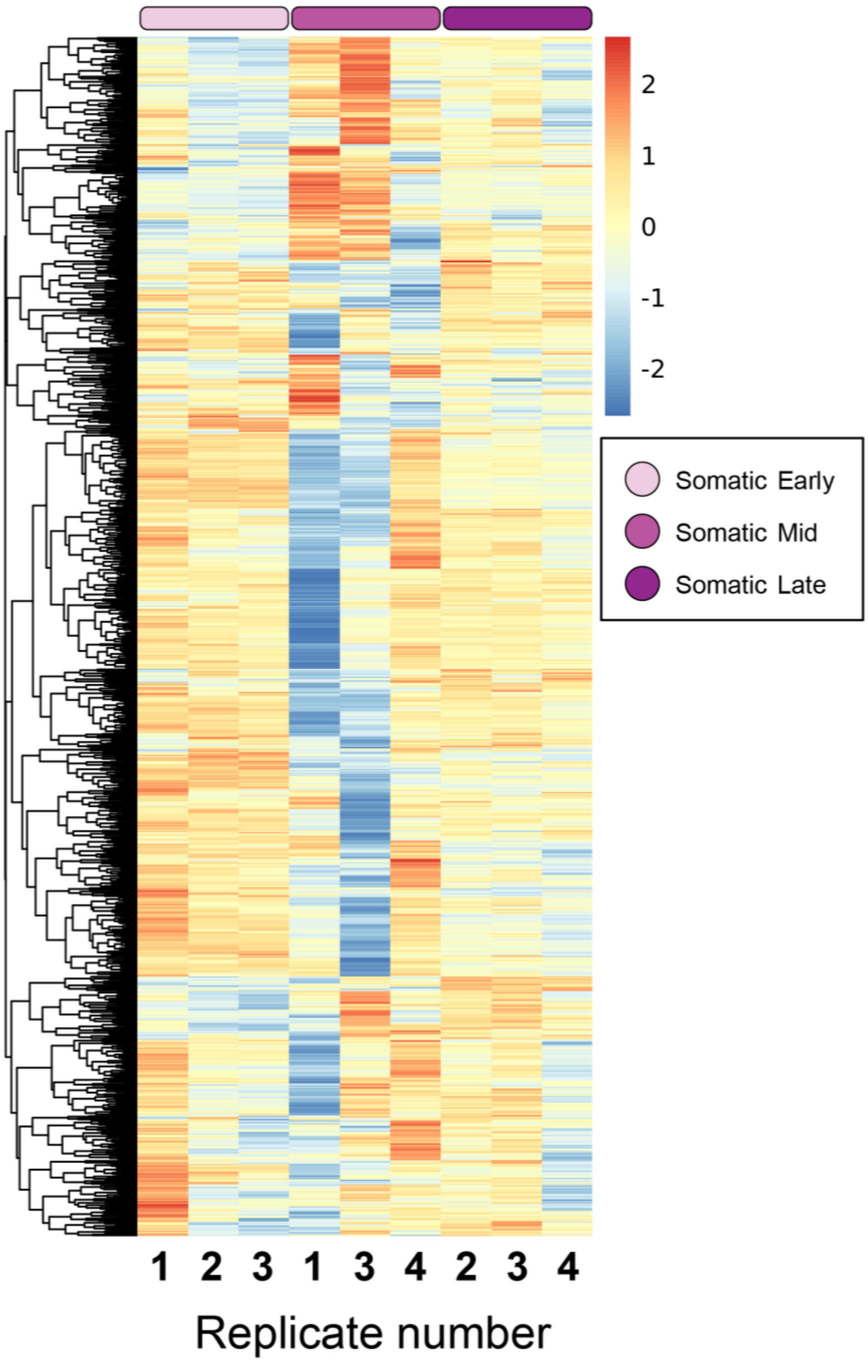


**Supplementary Figure S12.** Expression across stages in germ cell libraries of the germ cell markers obtained from SLAIDINA *et al.* (2020). Genes are grouped based on hierarchical clustering. The expression of each gene across samples is represented as a row-wise Z-Score value of the VST-normalized counts from high (red) to low (blue).

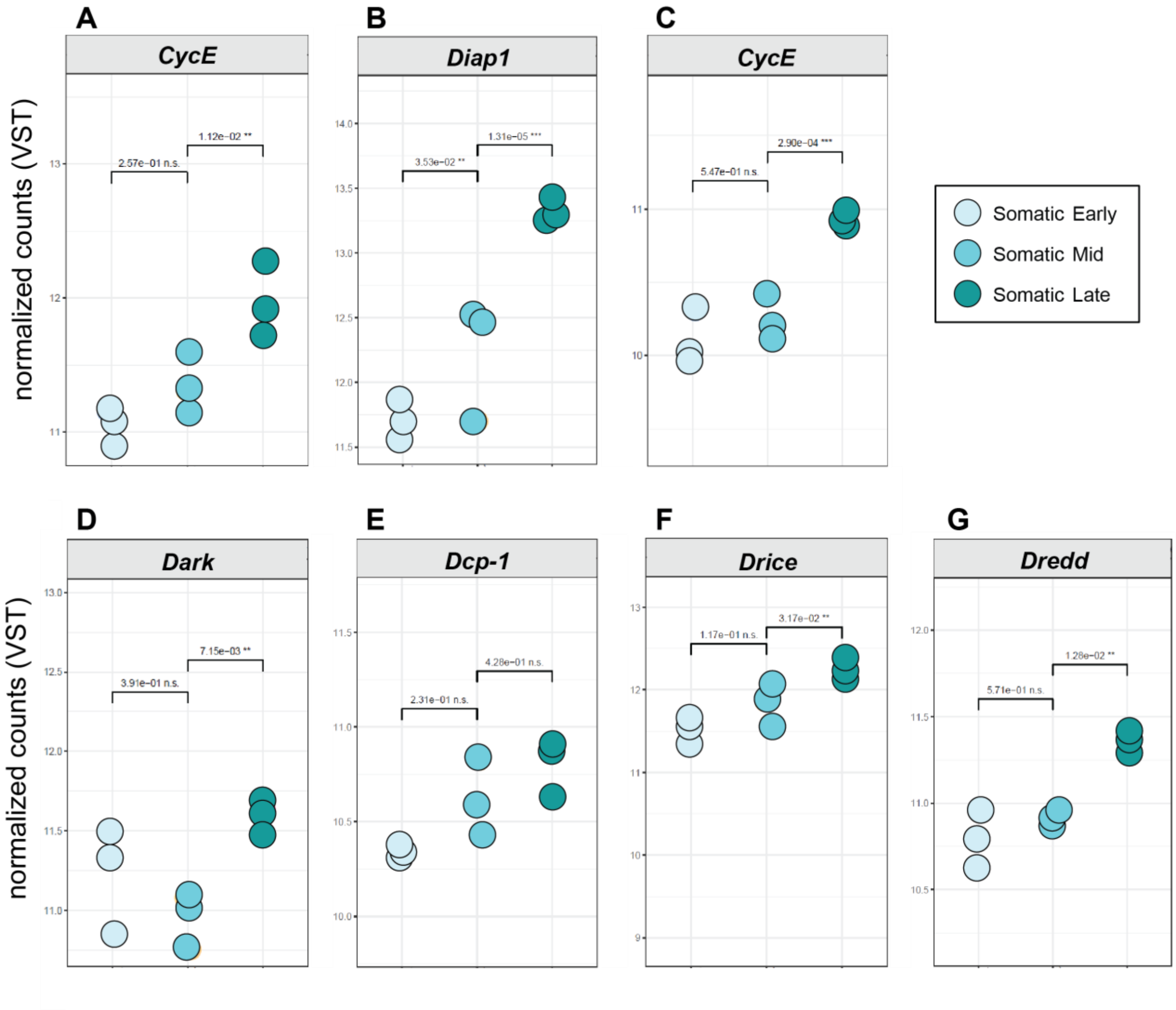


**Supplementary Figure S13** Dot plots of selected proliferation and apoptosis control genes differentially expressed across stages. Expression in VST counts of genes in each tissue-specific RNA-seq library.
